# Measuring linkage disequilibrium and improvement of pruning and clumping in structured populations

**DOI:** 10.1101/2024.05.02.592187

**Authors:** Ulises Bercovich, Malthe Sebro Rasmussen, Zilong Li, Carsten Wiuf, Anders Albrechtsen

## Abstract

Standard measures of linkage disequilibrium (LD) are affected by admixture and population structure, such that loci that are not in LD within each ancestral population appear linked when considered jointly. The influence of population structure on LD can cause problems for downstream analysis methods, in particular those that rely on LD pruning or clumping. To address this issue, we propose a measure of LD that accommodates population structure using the top inferred principal components. We estimate LD from the correlation of genotype residuals and prove that this LD measure remains unaffected by population structure when analyzing multiple populations jointly, even with admixed individuals. Based on this adjusted measure of LD, we can perform LD pruning to remove the correlation between markers for downstream analysis. Traditional LD pruning is more likely to remove markers with high differences in allele frequencies between populations, which biases measures for genetic differentiation and removes markers that are not in LD in the ancestral populations. Using data from moderately differentiated human populations and highly differentiated giraffe populations we show that traditional LD pruning biases *F*_ST_ and PCA but that this can be alleviated with the adjusted LD measure. In addition, we show the adjusted LD leads to better PCA when pruning and that LD clumping retains more sites and the retained sites have stronger associations.

## Introduction

Linkage disequilibrium (LD) is a measure of non-random association between alleles at different sites. If the frequency of a haplotype carrying a particular pair of alleles is equal to the product of the frequencies of the respective alleles, then the alleles are independent and are said to be in linkage equilibrium; otherwise, there is some degree of LD between the two alleles.

Drift and mutation will cause LD, whereas recombination tends to break it down. Therefore, alleles at sites located close to each other on the genome are likely to be in high LD since recombination events between those sites are rare. Conversely, alleles at sites located far apart or on different chromosomes generally have lower levels of LD. However, various other biological processes create and maintain LD, including selection, inbreeding, and population structure [1]. In this study, we focus on the latter, where differences in allele frequencies among populations cause LD when two or more populations mix or are considered jointly [2, 3]. LD created in this way may be observed at long genetic distances, including between chromosomes.

For many purposes, the presence of LD can be very useful. LD is at the heart of genome-wide association studies (GWAS), where the presence of non-genotyped causal SNPs might be detected from LD with a subset of genotyped SNPs, using a SNP-chip [4]. Furthermore, the overall pattern of LD itself constitutes a signal on which population genetic inference may be based, since the way LD decays as a function of distance can be used to estimate the effective population size through time [5–7] and the timing of admixture events [8, 9].

On the other hand, the presence of LD can equally present a challenge. In the GWAS setting, multiple SNPs will typically be in LD with a particular causal SNP, leading to spurious duplication of associations, which makes it difficult to determine the causal SNPs and the corresponding genes. Moreover, a wide variety of analyses in population genetics assume that sites are independent, and this assumption is clearly violated in the presence of LD. Therefore, LD may bias results in the context of admixture inference [10] or for detecting population structure using PCA [10, 11]. Various methods exist to address this issue, the most popular of which is LD pruning, where SNPs are removed in a way to ensure that all SNP pairs within a certain distance have LD below a predefined threshold.

Unfortunately, LD pruning can itself be problematic in some contexts. Standard LD pruning methods calculate the genotype correlation between two SNPs and remove the one with the lowest frequency if they are in LD (e.g. *r*^2^ greater than some cutoff). However, if the sample consists of individuals from multiple populations, SNPs with larger differences in sub-population allele frequencies will be more likely to be pruned due to the correlation introduced by population structure. This is sometimes referred to as the two-locus Wahlund effect [2, 12, 13]. The consequence is that SNPs with high genetic differentiation between populations are more likely to be removed by standard LD pruning, which causes populations to appear more similar than they are. This has been shown to bias *F*_ST_ [14], but also affects a range of other methods that quantify genetic difference, as we show in this study.

We address this problem by using a measure of LD which adjusts for the effects of population structure and admixture. We show that PCA can be used to model individual allele frequencies, predict individual genotypes, and then use the residual differences between observed and predicted genotypes to calculate LD. Using real data from populations with moderate differentiation such as humans and high differentiation such as giraffes, we show that standard LD is inflated by population structure, which causes biases in downstream analyses. We demonstrate that our proposed adjusted LD measure greatly reduces these biases.

## Methods

### Standard LD measures

For measuring LD between SNPs in a homogeneous population, standard statistics are based on haplotype frequencies [15]. Given two diallelic SNPs at position *s* and *t*, the (theoretical or true) haplotype covariance is

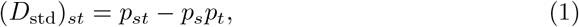

where *p*_*st*_ is the probability of having both reference alleles at positions *s* and *t*, and *p*_*s*_ and *p*_*t*_ are the probabilities of having the reference allele at *s* and *t*, respectively. This measure is then used to define the haplotype squared correlation as

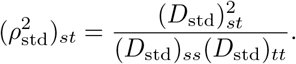

These two quantities might then be estimated using empirical haplotype frequencies [16]. However, both the theoretical quantities and the estimates fall short if the population is not homogeneous but has structure.

### LD in admixed populations

For admixed populations, we are interested in a measure of LD that takes into account the genetic heterogeneity between sub-populations. The main difference from the homogeneous case is that the parameters describing an individual genetic composition are now in part private to the individual, and depend on the specific admixture proportions of the individual, that is, *D*_std_ and *ρ*_std_ are no longer meaningful quantities.

We propose an ancestry adjusted measure of LD between genomic sites. Since genomic data are generally unphased, the measure is defined from genotype data rather than haplotype data. We first present a theoretical measure of true LD and afterwards provide means to estimate this measure from observed genotypes.

### Sample LD

Let *n* be the sample size and *G*_*is*_, *G*_*it*_ ∈ {0, 1, 2}, *i* = 1, …, *n*, be the number of reference alleles in two sites, *s* and *t*, respectively. Furthermore, let 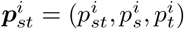 be the parameters of individual *i* = 1, …, *n*, where 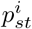 is the probability that a haplotype carries both reference alleles, and similarly for 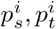. Using that the number of reference alleles in each site is a sum of independent parental gametic contributions 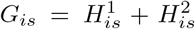 and 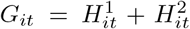. If we assume that both haplotypes are independent and identically distributed, the expression for the covariance between the genotype numbers is

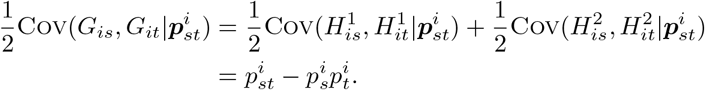

A measure of the (true) sample LD is thus the average covariance over all *n* individuals,

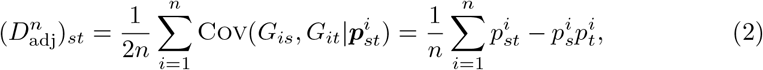

which is adjusted for the heterogeneity in the sample. Clearly,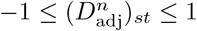. If the parental haplotypes do not share the same distribution, then (2) becomes a sum over the 2*n* haplotypes. Moreover, if there are *k* separate sub-populations and *n*_*ℓ*_ individuals from the *ℓ*th sub-population, our measure of population LD agrees with that of [2],

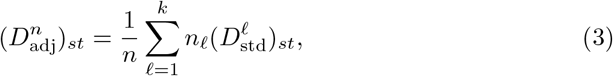

where 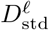 is the standard measure of the *ℓ*th sub-population. Hence, the proposed measure of population LD also extends previous work on LD in sub-divided populations.

An adjusted squared correlation might be defined similarly to the standard haplotype squared correlation. If the tuple 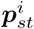 is the same for all the individuals, then 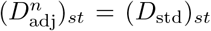, which shows that the adjusted measure of sample LD and the standard measure of LD agrees in the case of a homogeneous population. In that case, there is no dependence on the sample size *n*.

### Population LD

It is desirable to have a measure of LD that reflects the admixed population as such and is not attached to a specific sample of individuals. One might imagine taking *n* → ∞ in (2) assuming that the parameters 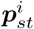 for each individual follow a common distribution ***P***_*st*_. It is thus natural to replace the average in (2) with an expectation. For this, let ***p***_*st*_ be a random draw from ***P***_*st*_, and let *G*_*s*_, *G*_*t*_ ∈ {0, 1, 2} be the number of reference alleles in the two sites, drawn according to ***p***_*st*_. Then, we define the (true) population LD as

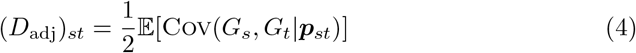

where the expectation is with respect to ***P***_*st*_. Also, the population LD might be written in terms of the haplotype covariance. It follows from the law of large numbers that 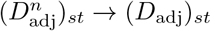 as *n* → ∞. An adjusted squared correlation might be defined similarly to the standard haplotype squared correlation.

If the distribution ***P***_*st*_ is degenerate (always takes the same value), which is the case if the population is homogeneous, then (*D*_adj_)_*st*_ = (*D*_std_)_*st*_. Thus, the proposed measure of population LD extends the standard measure of LD. In the case of *k* separate sub-populations with relative population sizes *w*^*ℓ*^, *ℓ* = 1, …, *k*, then the population LD is a sum over the standard LDs of the sub-populations, similarly to for the sample LD,

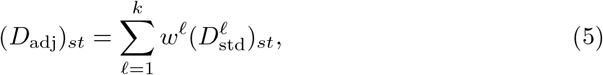

where 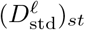 is the standard LD measure of the *ℓ*-th sub-population. In fact, a stronger result holds.

**Theorem 1**. *Let*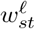, *ℓ* = 1, …, *k, be the probability that the two alleles of a haplotype are both from sub-population ℓ. Then*, 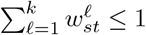, *and*

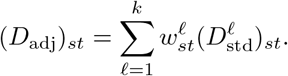

*Furthermore, by Jensen*’*s inequality*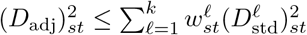.

In particular, (*D*_adj_)_*st*_ always takes a value between the smallest and the largest value of 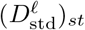, *ℓ* = 1, …, *k*. Similarly, 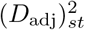 is always smaller than the largest value of 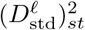, *ℓ* = 1, …, *k*. In contrast, there is no lower bound but 0. For example, in the case where there is a balanced pooling of two sub-populations with 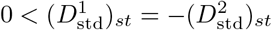. Then,

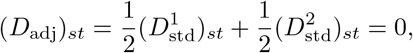

and hence 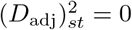, even though 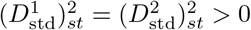.

### Estimation of LD in admixed populations

To estimate the sample LD, we first introduce a model and some notation. Let *G* be the observed genotype data matrix of *n* individuals across *m* SNPs, where each genotype consists of two alleles, hence each entry of *G* is 0, 1, or 2. Let Π_*is*_ ∈ [0, 1] be the probability that individual *i* has the reference allele at position *s*, so both *G* and Π are matrices of dimension *n×m*. We model the marginal distribution of *G*_*is*_ as a binomial distribution

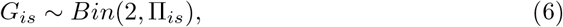

and allow for dependence between *G*_*is*_ and *G*_*it*_ (within one individual), but we do not specify a model of this explicitly. In contrast, we assume genotypes from different individuals are independent, that is, *G*_*is*_ and *G*_*jt*_ are independent for *i* ≠ *j*, given Π_*is*_ and Π_*jt*_. In the context of the previous section, 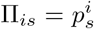, and 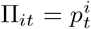, whereas we leave unspecified the form of 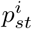.

Based on this general model, we introduce an estimate of the ancestry adjusted sample LD by first calculating the empirical covariance of the *n × m* residual matrix,

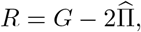

where 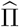 is a matrix of predicted (estimated) values of Π. In this way, we purge the component of covariance generated by population structure, while keeping the linkage between SNPs. Then, the empirical covariance between two columns of *R* using the Bessel’s correction is an estimate of the ancestry adjusted sample LD,

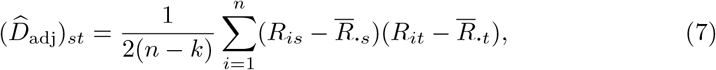

Where 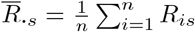, and *k* is the rank of Π. The empirical squared correlation, an estimate of the adjusted squared correlation, is given by Pearson’s correlation calculated on the residuals, that is

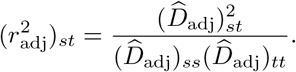

The estimator in (7) is applicable whenever Π is estimable. We estimate Π assuming it has a specific structure, namely that the ancestry of each individual is composed of genetic material from *k* ancestral populations, where we take *k* to be a known parameter. Specifically, we assume that the matrix Π factorizes as Π = *QF*, where *Q* is an *n × k* matrix of rank *k* ≤ *n* consisting of (true) ancestral admixture proportions, such that the proportion of individual *i*’s genome from population *ℓ* is *Q*_*iℓ*_ with 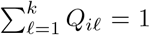 and *F* is an *k × m* matrix of (true) ancestral SNPs frequencies, such that the frequency of the reference allele of SNP *s* in the ancestral population *ℓ* is *F*_*ℓs*_. Hence, the probability that an individual *i* has the reference allele in site *s* is 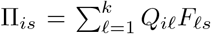. This is similar to admixture models proposed in the literature [17, 18] and models based on PCA [19, 20]. The rank condition on *Q* is for reasons of identifiability, to be able to disentangle the admixture proportions from all *k* ancestral populations for each individual.

To estimate Π = *QF*, different methods might be used. We distinguish between whether *Q* is known (e.g., when the population is homogeneous, *k* = 1 and *Q* is a vector of length *n* with only ones) or unknown. In the former case, we use linear regression to obtain an estimate of 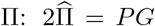, where *P* = *Q*(*Q*^t^*Q*)^−1^*Q*^t^ is an *n ×n* matrix of rank *k*, which is the projection onto the column space of *Q*. Here, *M*^T^ denotes the transpose of a matrix *M*. If *Q* is unknown, one might use PCA to estimate Π by projecting *G* onto the first *k* principal components [19–22]. If so, then 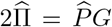, where 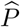 is an estimate of *P* [23]. Alternatively, one might estimate Π by first estimating *Q* and *F*, as suggested in [17, 18]. In that case, an estimate of Π can be obtained either as 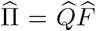 or as 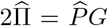, where 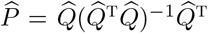 is an estimate of *P*.

Whenever an estimate 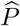 of *P* is available, we have

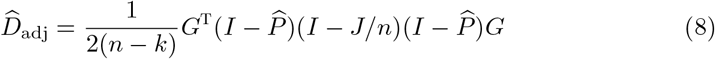

(potentially with 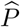 replaced by *P*, if *Q* is known), where *J* is an *n × n* matrix with all entries equal to one, see Lemma S1. In the case *Q* is known, (8) reduces to

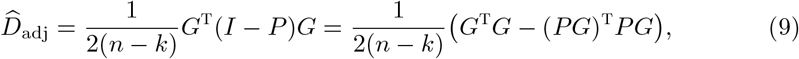

using that *P* ^2^ = *P* for a projection matrix. Since *PG* is an estimator of the expectation of *G*, then the second expression of 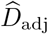 above resembles that of 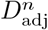.

If the population is homogeneous (*k* = 1 and *Q* is a vector of ones), then 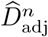 agrees with the unbiased estimator of *D*_std_, based on unphased genotype data, suggested by [24] (there is a factor 2 missing in their expression for 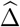 on p931), and Burrow’s estimator of *D*_std_, also based on unphased genotype data [16, 25]. Hence, the estimator 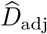 is an extension of known estimators for LD. Similarly, in the case of a sub-divided population into *k* separate sub-populations with *n*_*ℓ*_ individuals from the *ℓ*-th sub-population, with *ℓ* = 1, …, *k*, and

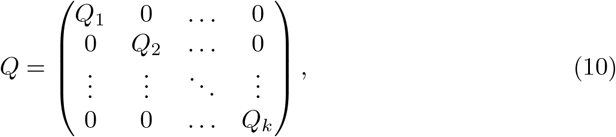

where *Q*_*ℓ*_ is a column vector of length *n*_*ℓ*_ with all ones, and the 0s are null vectors of matching lengths, then 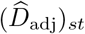 agrees with the estimator suggested in [2], when adapted to unphased genotype data using Burrow’s estimator.

### **Properties of** 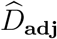

It remains to connect the estimated ancestry adjusted LD, 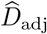, to the true population LD, *D*_adj_. In the following, we add superscripts *n, m* to emphasize the dependence on *n, m*, the size of the data matrix.

We explore the case where the sample size *n* and the matrix *Q* are fixed (none random), even though the admixture proportions might be known or unknown. In that case, we simply write 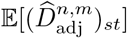 for the expectation of 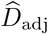. On the contrary, when *n* is large, that is, when *n*→∞, we consider the admixture proportions of each individual as a random draw from the distribution given by the population.

**Theorem 2**. *Assume* 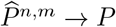*as m* → ∞ *with n fixed. Then, it holds that*

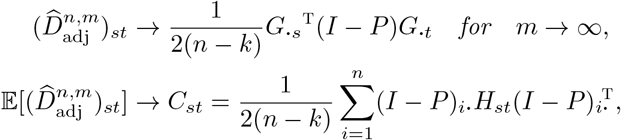

*where H*_*st*_ *is the n × n diagonal matrix with diagonal elements* Cov(*G*_1*s*_, *G*_1*t*_), …, Cov(*G*_*ns*_, *G*_*nt*_), *the LD between the two SNPs for each individual*.

*In particular, if G*_***·****s*_ *and G*_***·****t*_ *are independent, then*

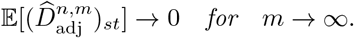

*Moreover, if Q is known, then* 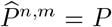 *and the above holds with* → *replaced by* = *without the need of m* → ∞.

Conditions for when 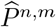 converges are given in [23]. In particular, the PCA approach suggested by Chen and Storey [21, 26] guarantees convergence. Under additional conditions, the same holds for standard PCA based on the mean normalized genotype data matrix [23]. Empirically, convergence seems to hold irrespective the PCA approach used, or whether some other method, for example [17], is applied to obtain 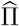 or 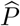. This is important because in practice on large data sets, the PCA approach of [21, 26] has severe computational limitations. In data analysis, we used mean normalized PCA, as this is conventionally used.

The limiting expression of 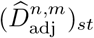 in Theorem 2 is identical to that of 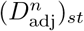 in (9) with the same interpretation. Moreover, in the case where we consider the pooling of *k* separate sub-populations, with *n*_*ℓ*_ individuals in sub-population *ℓ* as in (10), *C*_*st*_ is the pooled covariance of each sub-population with the Bessel’s correction (see Lemma S2), that is,

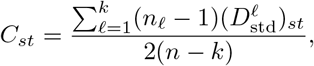

which we can compare with (5). In particular, *C*_*st*_ agrees with the sample LD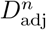, when there is only one ancestral population.

**Theorem 3**. *If Q is known, then for any pair of SNPs s and t*,

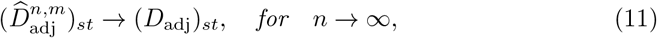

*the population LD*.

When *Q* is known, as in Theorem 3, then 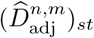 does not depend on *m*.

### **Sample size correction for mean** *r*^2^

When calculating the mean squared correlation coefficient the sample size becomes important because this measure is biased for finite sample sizes. There are several suggested methods for correcting this bias, but none of them perform perfectly [24]. We choose to use the method used in LD score regression [27] due to its simplicity. The correction is given by

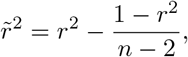

where *r*^2^ is the calculated squared correlation coefficient and 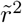 is the bias-corrected. However, it should be noted that it is not trivial to correct for this bias [24] and that other methods also exist that perform similarly well [24, 28] in mitigating the upward bias.

## Results

In the following sections, we compare adjusted LD to standard LD on real data. We are interested in the measures themselves as well as their effects on downstream analyses when used for pruning and clumping.

### Data

To illustrate the problems with standard LD in the presence of population structure, we use two datasets: one with moderately differentiated populations and one with a large amount of differentiation. In both cases, we have high quality SNP and genotype calls from medium or high depth whole genome sequencing data so that no prior SNP ascertainment was done other than quality control. An overview is shown in fig. 1a.

**Figure 1.**
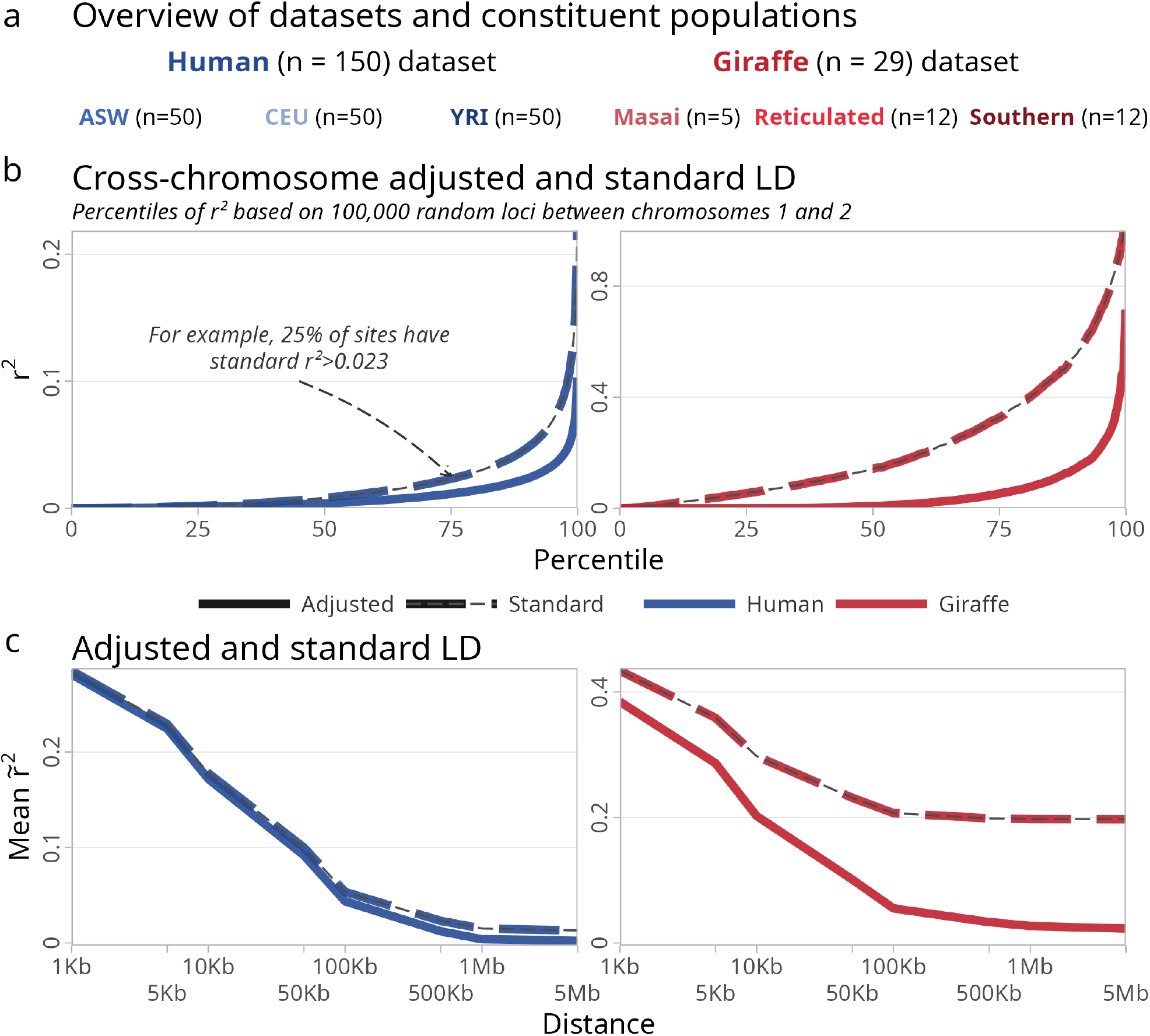
Comparison of LD measures. a: An overview of the two datasets and the sub-populations included within them. b: Standard and adjusted *r*^2^ based on 100 000 randomly sampled cross-chromosome pairs of sites. Percentiles of observed *r*^2^ based on these samples are shown. c: LD decay curves of standard and adjusted 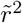. Mean 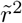 shown in bins based on a sliding window over sites up to 5 Mb apart.

First, for the case of moderate population structure, we use the high-quality, human data from the 1000 Genomes Project [29]. Specifically, we used 50 random, unrelated individuals from each of the CEU (Utah residents with Northern and Western European ancestry), YRI (Yoruba in Ibadan, Nigeria), and ASW (African ancestry in Southwest US) populations. The latter was chosen because the African Americans represent an admixed population with European and East African ancestry. We used PLINK [30, 31] to subset the data and remove sites with minor allele frequency (MAF) less than 5 %. The resulting dataset contains approximately 10 M common variants.

Second, we also included a dataset of whole-genome sequencing (approx. 20 x coverage) of three populations of giraffes [32, 33], which we refer to as Masai (*n* = 5), Reticulated (*n* = 12), and Southern (*n* = 12). In comparison to the human dataset, the sample sizes are lower, and the population structure significantly greater—indeed, these groups might be considered different species [32], though we use the term populations throughout.

### Measures of LD

There are many ways to calculate LD [24]. We choose to focus on the squared correlation coefficient as this is an often used measure and because it is used in LD pruning. We used the *r*^2^ obtained directly from the covariance matrix and when adjusting the *r*^2^, we used PCA based on mean centered genotypes. However, other approaches for PCA and *r*^2^ might be used, but in our analysis, they performed similarly on the above data sets (see fig. S14 and fig. S15).

We begin by comparing 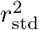 and 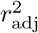 LD between variants on different chromosomes. As described, standard measures of LD, including 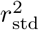, are expected to find LD between sites on separate chromosomes in the presence of population structure, despite little or no LD being present in the ancestral populations prior to mixing. In contrast, we expect 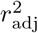 to be generally close to zero in the cross-chromosome case, since it adjusts for this effect.

To investigate, for each dataset we randomly sampled 100 000 pairs of SNPs split between chromosomes 1 and 2. For each of these pairs, we calculated 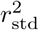 and 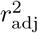. The results (fig. 1b) confirm that the adjusted measure significantly reduces the amount of cross-chromosome LD measured. For example, on the giraffe dataset, the median 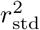 is 0.14, whereas for the adjusted measure, the median 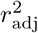 is 0.008. As expected, the difference between adjusted and standard LD is greater on the giraffe dataset with greater population differentiation, but it is likewise apparent in the human data where 9.1 % and 1.6 % of SNP pairs have *r*^2^ *>* 0.05 for standard and adjusted LD, respectively.

We then turn to demonstrate that adjusted LD is meaningful within chromosomes, and not just a decreased measure of LD. When there is LD in the ancestral population, we expect both methods to capture this, but we expect standard LD to plateau to a higher level with increasing distance, since the relative importance of population structure on 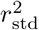 is larger at longer ranges. This can be seen in the LD decay curves shown in fig. 1c for both standard and adjusted 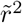 up to distances of 5 Mb. (The same data is shown without sample size correction in fig. S1. Standard LD curves for the separate populations are shown in figs. S2 and S3, with SNPs private to each population having been removed.) On the giraffe data, the standard LD curve is significantly shifted up relative to the adjusted measure. Again, the effect is less visually apparent on the human dataset, but we note that although smaller, the difference is due to sites that are particularly informative of population structure and hence are expected to exert an outsize influence on certain analyses. We return to this point when looking at pruning below.

### Effects of pruning

The LD measure has an effect on LD pruning and analyses based on pruned data. To investigate, we implemented a pruning algorithm like the one used in PLINK [30].

Briefly, for each SNP A, we consider all SNPs B in a window up to 100 kb ahead. For each SNP B in the window, starting with the closest, we calculate *r*^2^ (either adjusted or standard) and remove the SNP with the lowest MAF if the value is above 0.5. The process is repeated until either the starting SNP A is removed in this way, or the end of the window is reached, and the window is then moved one SNP forward. Pruning occurs separately for each chromosome.

We pruned both datasets using either 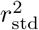 or 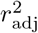. To see the direct effects of pruning, we then calculated the standard LD decay curve from the jointly pruned data. In addition, we extracted genotypes from the jointly pruned data and calculated the standard LD curve for the common variants for each of the three populations. The resulting LD curves (fig. 2a, without sample size correction in fig. S4) show that using 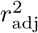 over 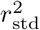 has a large effect on the joint LD curve (where there is population structure), but a comparatively smaller effect on the curves when calculating the remaining LD for each separate population. In other words, where there is population structure, pruning based on 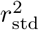 removes more LD (by standard measures) than 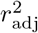 by removing sites in LD due to population differentiation while both methods remove a similar amount of within population LD.

**Figure 2.**
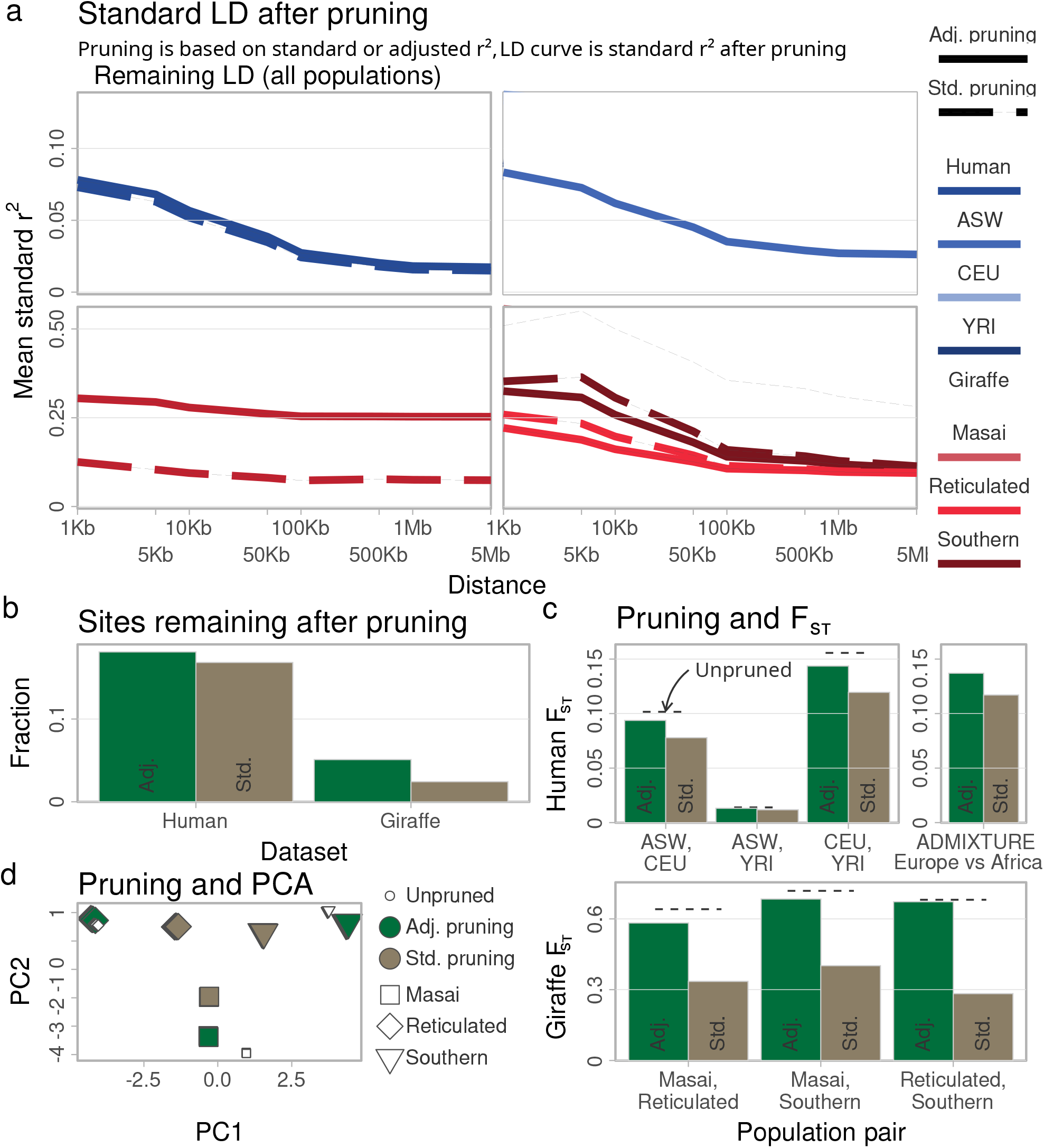
Effects of pruning using different LD measures. a: Standard LD decay curves after pruning based on either standard or adjusted *r*^2^. The joint datasets are shown, as well as LD decay curves for the constituent populations extracted from the pruned datasets. b: Fraction of sites remaining after pruning. c: *F*_ST_ after pruning for each pair of populations, with the unpruned *F*_ST_ shown for comparison. d: PCA after pruning on the giraffe dataset, with the unpruned data included for comparison. Eigenvectors scaled by corresponding eigenvalue shown.

As a consequence of removing sites in LD due to population structure, standard pruning also removes more sites than adjusted pruning (fig. 2b). On the giraffe dataset, more than twice as many sites are retained when pruning based on 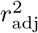 compared to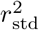. Again, the differences on the human dataset look less stark. For example, only about 10 % more sites are kept after pruning by 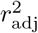 on the human dataset. However, those extra sites are likely to be exactly those that are most informative of population structure. As a result, our ability to infer population structure will be diminished by standard pruning. To illustrate, we used PLINK2 [31] to compute Hudson’s estimator of *F*_ST_ [34, 35] for all population pairs before and after pruning. The results, in fig. 2c, show significant differences in the estimates. For example, pruning based on standard and adjusted *r*^2^ lead to *F*_ST_ estimates for CEU and YRI of 0.119 and 0.144, respectively, compared to a value of 0.156 using the unpruned data. On the giraffe dataset, the deviation is more extreme, with *F*_ST_ values after standard pruning approximately half those resulting from adjusted pruning.

As can be seen, *F*_ST_ remains lower after pruning even when using the adjusted measure. A possible explanation, which we ruled out, is that this is due to the direct effects of pruning on the one-dimensional frequency spectrum, shown in fig. S5 before and after pruning. To test this, we re-sampled sites after both standard and adjusted pruning in frequency bins of 0.001 to match the unpruned frequency spectrum (fig. S6). Re-calculating *F*_ST_ on these re-sampled datasets results in broadly similar patterns (fig. S7). We note that LD pruning is not typically required for standard *F*_ST_ calculations, so these results mainly serve to illustrate the differences between the pruning methods in the context of inferring population structure. However, even for *F*_ST_, the effects of LD pruning are sometimes important. For instance, if we wish to infer the *F*_ST_ of the ancestral components as inferred e.g. by `ADMIXTURE` [36] software, pruning is assumed by the standard admixture model. To illustrate the effect of the LD adjustment in this context, we ran `ADMIXTURE` for 10 different seeds on each of the two pruned datasets, and recorded the *F*_ST_ value for the run with the highest log-likelihood. The results are included in fig. 2c and show a similar reduction of *F*_ST_ when using standard LD pruning.

Based on the differences in *F*_ST_ and the relationship between *F*_ST_ and PCA [37], it is expected that a PCA to be likewise affected by the pruning method. Running PCA in PLINK confirms that this is the case on the giraffe data fig. 2d. As expected from the *F*_ST_ results, both methods of pruning affect the shape of the principal components, though the adjusted LD pruning has a much smaller impact. The human data (fig. S8) shows the same pattern along the first component, but is harder to interpret along the second, since there is only one main axis of variation relating to population structure in the human dataset. In addition, the small size of both the giraffe and human datasets makes it hard to judge whether the eigenvectors themselves are impacted, or whether the pruning LD measure only influences the scaling.

### PCA

To explore this question, we analyze the use of adjusted pruning for PCA on a larger and more complex human dataset. Specifically, based on the 1000 genomes dataset without close relatives (first and second degree) [29], we carried out both standard and adjusted pruning on the common variants (MAF*>* 5%). We performed PCA on each of the resulting data sets, as well as on the unpruned data for comparison. We chose to adjust the adjusted LD with the top 8 PCs from unpruned data since these capture population structure (see fig. S9).

The top four principal components are shown in figs. 3a and 3b, comparing the two pruning methods to the unpruned PCs. The other top PCs are shown in fig. S9 and fig. S10 which also shows that our standard pruning algorithm is comparable to the one in PLINK [30, 31]. Looking at the raw eigenvectors unscaled by the eigenvalues, we do see subtle differences between the pruning methods. Moreover, as we move to the higher PCs, we begin to see that some of the PCs are shuffled, i.e. that the different axes of variation are captured in different order. The unpruned data also starts to capture LD as indicated by locally high SNP loading. The question remains, however, whether these differences are meaningful and, if so, which method is preferable.

**Figure 3.**
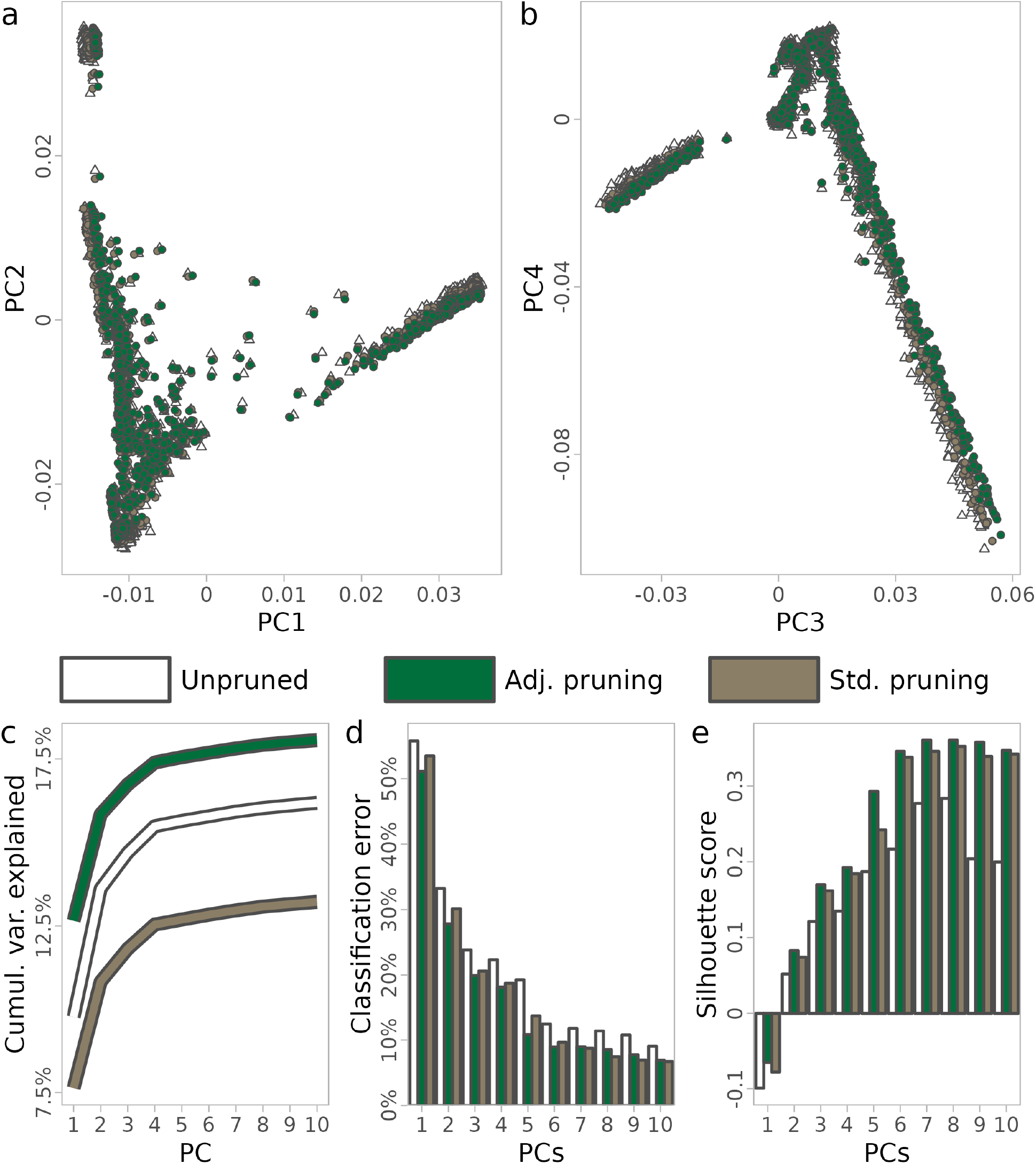
PCA results on full 1000G data set comparing no pruning, and pruning based on either standard or adjusted LD in a window of 1000 kb with a *r*^2^ cutoff of 0.2. a, b: First four principal components, *not* scaled by eigenvalues. Population labels can be seen in fig. S9 and fig. S10. c: The cumulative variance explained for the first ten components. d: The classification error for clustering using `mclust` with the population labels. c: The mean silhouette score for clustering based on population labels.

As a first answer to this question, fig. 3c compares the cumulative variance explained for the first 10 PCs and clearly shows that PCA based on adjusted pruning explains more of the variance in the genotype data using fewer components. While informative of the structure of the decomposition overall, the variance explained measure ignores our knowledge of the population labels. In other words, it does not quantify how well each method captures population structure specifically. To investigate this, we examined specifically the separability and clusterability of the 26 populations in the PCAs, taking the given population labels from the 1000G project as ground truth. We took two distinct approaches to this.

First, we used the `mclust` R package [38] to cluster the populations using a variable number of the top PCs. For this, `mclust` does an automated model selection of different parametrizations of Gaussian mixtures to perform supervised clustering into the 23 population labels. Looking at the resulting classification errors (fig. 3d), we see that adjusted pruning leads to lower classification errors for the first 5 PCs, after which the difference between adjusted and standard pruning tails off. While small, the benefit of adjusted pruning is consistently on the order of a couple of percentage points, which is a meaningful difference. The corresponding Brier scores are shown in fig. S11 and show a similar pattern.

Second, we use `scikit-learn` [39] to compute silhouette scores [40] as a measure of clusterability by comparing distances between individuals from the same or different populations. As above, we varied the number of top PCs and averaged the silhouette score across all individuals, with results shown in fig. 3e (Figure S12 shows the data broken down by population). Again, the results indicate a consistent benefit of using adjusted LD pruning for PCA analysis compared to both no pruning and standard LD pruning.

### Clumping

A final application for which the choice of LD measure matters is in the context of association studies where clumping is performed to obtain association signals that are independent from each other. This is a standard practice when for example performing Mendelian randomization[41]. While good methods exist for accounting for the effects of population structure while estimating the associations themselves (e.g. mixed models [42]), these do not extend to the process of identifying unlinked loci afterwards. Briefly, clumping is one solution to this problem in which, starting from the most significant variants, all other variants within some distance are removed if they are in LD above some threshold, after which the procedure is iteratively applied to the next most significant variant that has not yet been removed. Clumping may be preferable to pruning for association studies, since clumping takes into account the inferred *p*-values to keep the most significant hit in each group of linked loci.

However, we anticipate that LD induced by population structure would serve to interfere with this process. Specifically, the idea is that clumping based on standard LD would discard SNPs that are not in LD when populations are considered separately and thus are associations that only appear correlated due to population structure. For most applications, the removal of such SNPs is undesired as the associated ones are needlessly removed and, as we show, because the retained SNPs have a weaker association to the traits.

To investigate, we applied clumping to summary statistics of a cross-population BMI GWAS study [43] (GWAS catalog [44] accession: GCST90018947). To perform clumping, we calculated standard and adjusted LD based on the 1000 genomes data. We ran the clumping algorithm with various cutoffs on *r*^2^ (0.0005, 0.001, 0.002, 0.005, 0.01, or 0.02) and various maximum distances within which SNPs can be removed (1 Mb, 5 Mb, or entire chromosomes). We refer to this procedure as adjusted or standard clumping, respectively, corresponding to the input LD measure. The *r*^2^ thresholds chosen are stringent to illustrate the performance for clumping for Mendelian randomization. For reference, the default *r*^2^ threshold used in the popular package `TwoSampleMR`[45] is 0.001.

We find that for each combination of LD cutoff and clumping distance, adjusted clumping retains at least as many, and often more, association hits as standard LD(fig. 4a). To quantify the strength of the association between these SNPs, fig. 4b shows the sum of χ^2^ scores for the kept SNPs, expressed as the difference between the two clumping methods. Across the range of LD cutoffs and clumping distances considered, this combined association strength of the kept SNPs is much higher using adjusted LD, suggesting that the quality of hits retained with adjusted clumping is preferable.

**Figure 4.**
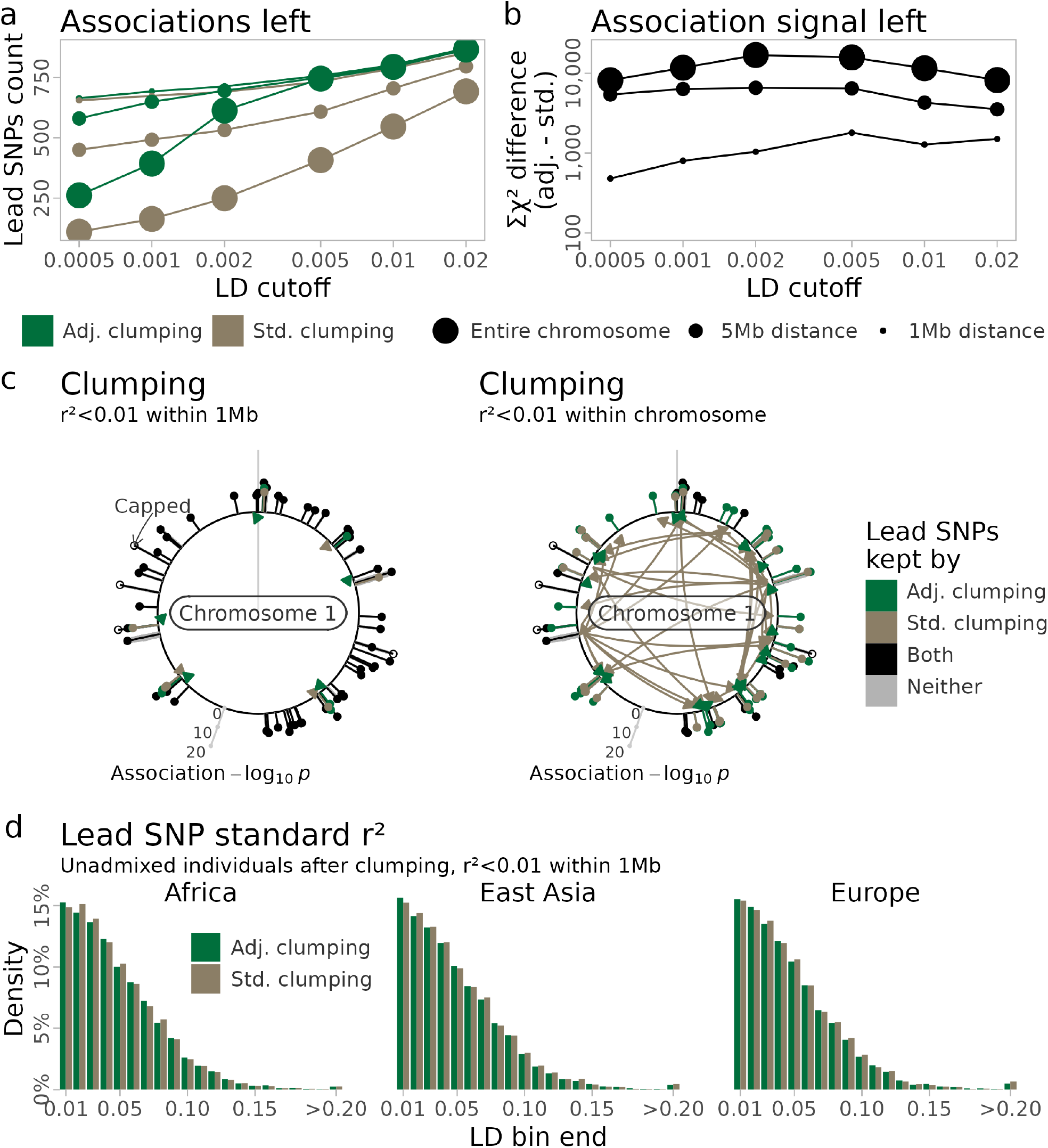
Summary of stats for clumping from a multiethnic GWAS study comparing adjusted and standard LD estimated from the 1000G data. a: Inferred lead SNPs left after LD clumping using various *r*^2^ thresholds and maximum pairwise distances. b: Association signal after clumping as quantified by the difference in the sum of the χ^2^ statistics of the inferred lead SNPs. c: Chromosome 1 association signals coloured by the LD measure that retains it when clumping. −log_10_ *p*− values are capped at 20 for open circles. Each SNP kept by exactly one method has an arrow showing from which SNP there was an association that caused the removal on the other method. E.g., a green pin shows a SNP kept only with adjusted LD and has a brown arrow from the SNP that removed it when using standard LD. d: Standard LD left after LD clumping among 390 individuals from each of the African, East Asian, and European populations.

To visualize why adjusted LD performs better, fig. 4c shows the clumped SNPs on chromosome 1 according to clumping methods at the 0.01 LD cutoff for the 1 Mb and entire chromosome cases (cf. fig. S13 for 5 Mb). Inspection of these figures supports the claim that where adjusted and standard clumping differ, adjusted clumping retains peaks with stronger associations in a particular group of linked signals. The case with no maximum distance clearly illustrates the explanation: during standard clumping, long range LD induced by population structure removes a large number of hits that are independent in each of the constituent populations. Of course, this happens to a lower degree with a short distance cap, but this is arguably an *ad hoc* solution to the problem that adjusted clumping addresses in a more principled way. Moreover, as the 1 Mb and 5 Mb cases show, setting a cap only partially addresses the problem. Even though the choice of distance seems arbitrary, it greatly influences the selection of which SNPs are kept with standard clumping; in comparison, adjusted clumping is fairly robust to the chosen cap (if any) within a reasonable band of LD cutoffs, as argued above.

Finally, we want to confirm that despite keeping more highly significant association hits, adjusted clumping removes at least as much LD as standard clumping in the constituent populations considered separately. To illustrate this, we extracted 390 individuals from Africa (ESN, GWD, MSL, YRI), Europe (CEU, GBR, IBS, TSI), and East Asia (CHB, CHS, JPT, KHV) and calculated standard *r*^2^ among all intrachromosome pairs kept by each of the two clumping methods in the 1 Mb and *r*^2^ *<* 0.01 scenario. The corresponding LD distributions are indeed very similar, as seen in fig. 4d.

## Discussion

In this study, we developed the mathematical foundation of a simple to use method that provides a measure of LD. This measure has some desirable properties when applied to datasets with individuals from multiple populations. Most importantly, this measure does not increase when individuals from different ancestries are analyzed jointly, unlike standard LD. We prove that for samples that come from a mixture of *k* ancestral populations, then the expected adjusted LD is zero (*D*_adj_ = 0) if the LD in each of the ancestral populations is also zero (*D*_std_ = 0 within ancestral populations). This is achieved by subtracting the predicted covariance given by the population structure from the standard covariance matrix of the genotypes. In practice, we estimate the predicted covariance given by the population structure inferred from the top principal components. We show that, even with a finite number of individuals, the measure is unbiased as the number of SNPs goes to infinity. The estimator is also consistent such that, if there is no LD in the ancestral populations, then each pairwise adjusted LD goes to zero as the number of individuals and SNPs goes to infinity. When there is LD in the ancestral populations, then the adjusted LD measure is also correlated if the genotype covariance is bigger than what is predicted from the population structure. In particular, for separate sub-populations, the expectation of the adjusted LD is the pooled covariance of the ancestral populations.

We evaluate the performance of the adjusted LD based on two data sets. A giraffe dataset consisting of a pooling of 3 populations with a large amount of differentiation and a dataset of 2 moderately differentiated human populations including individuals that are a mixture of the two. To evaluate the measures when there is little ancestral LD we analyzed pairs of SNPs from different chromosomes. As expected the standard LD is high for many pairs of SNPs while it stays much closer to zero for the adjusted LD measure.

It is well known that analyzing individuals from multiple populations jointly increases standard LD. Hence, highly differentiated populations such as the giraffes, have a bigger increase in standard LD compared to more similar populations like humans. The difference is also observed on the LD decay curves, where the adjusted LD curves go towards zero as the distance between markers increases, while the standard LD curves levels at a much higher value. Because the mean estimated *r*^2^ is biased upwards for finite sample sizes, we show results for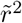, which is a simple correction based on the number of sampled individuals [27]. Other methods exist that in some instances perform better [24, 28]. However, we choose the simple correction due to its simplicity, but while taking into account that standard LD decay curves for the populations considered separately do not always approach 1*/*(*n* − 1) as predicted [27].

Our method is not the first method that tries to overcome the issues of population structure when calculating LD. A previous study [46] suggested first inferring the admixture proportions using *STRUCTURE* [18], and then using these as predictors in a linear model. From the linear regression, they obtain genotype residuals from which they calculate the correlation of residuals. This approach estimates the partial correlation [47] and assumes that the population structure is an observed variable.

This makes the approach prone to noise in the estimation of the confounding variable. Also, the confounding variable must be linear. That is not so in our case, where we assume that population structure has been estimated.

Measuring LD is of interest in itself, but it is also often used to make datasets more appropriate for further analysis. Many methods assume that sites are independent such that there is no LD in the ancestral populations. This includes commonly used methods for inferring population structure such as *ADMIXTURE* [17], *STRUCTURE* [18] and PCA [48]. However, it also includes frequent measures in population genetics such as *F*_ST_, *D*_*xy*_, heterozygosity, and kinship coefficients, which are often calculated under the assumption of no LD. Often it is not a big issue for the point estimates, since the correlation from LD mostly affects markers located close to each other. Thus the estimators can still be consistent [49]. However, the uncertainty of any estimates increases and, therefore, it is often recommended to perform LD pruning. This is not an issue for most analyses that are performed within a single panmictic population. However, if the samples come from multiple ancestral populations then standard LD pruning can cause biases[14, 50]. This is because LD is created between alleles with different ancestral allele frequencies: the so-called two-locus Wahlund effect [2, 12, 13]. Sites with a large difference in allele frequency are more likely to be pruned away because their standard LD increases more than sites with a small difference in allele frequency. Therefore, populations look genetically more similar after standard LD pruning. We illustrate this issue by performing LD pruning on the human and giraffe populations using both the standard LD measure and the adjusted one. The standard LD pruning removed slightly more sites than the adjusted in the human populations, while in the more differentiated giraffe populations, the adjusted pruning retained more than twice the number of SNPs compared to the standard LD pruning. However, if we calculate standard LD in the ancestral populations on the remaining sites then we see that both methods have similar amounts of standard LD after pruning. Thus both methods are able to greatly reduce the amount of ancestral LD, but the adjusted LD pruning can do it while retaining more SNPs. When calculating *F*_ST_ from the pruned data, we see that standard LD pruning causes a huge bias for the giraffe with *F*_*st*_ values being half of the value of the unpruned data. The effect is also apparent in the human where *F*_*st*_ is reduced by about 20%. Using the adjusted LD for pruning alleviates most of this bias but there still appears to be some negative bias left with *F*_ST_ values being around 5% lower than with the unpruned data. This remaining bias could be due to allele frequency ascertainment bias which is known to bias *F*_ST_ [51] and PCA. However, even if we sub-sample the pruned sites to match the overall allele frequency distribution (fig. S5), the bias remained (fig. S7).

In addition to *F*_ST_, we also explored the effect on PCA analysis, where standard pruned data showed fewer genetic differences between populations. This is not surprising since the top eigenvalues are proportional with *F*_ST_ [37] when analysing 3 populations. Nevertheless, the shape of the PCA was not affected for either the humans or the giraffes so in these cases the interpretation from the PCA would have remained the same.

Finally, we explored the performance of LD pruning and clumping on the diverse 1000 genomes project with individuals from 23 populations. We show that the PCA is better at reflecting population structure if the data is LD pruned, and that adjusted LD pruning performed better than standard LD pruning. The difference was very pronounced in the variance explained by each PC, but we also observed better performance in the clusterbility of the populations. To illustrate the advantage of adjusted LD clumping, we applied it to GWAS summary statistics from an ethnically diverse study. Standard LD clumping on this data set removed independent association signals caused by the population structure. This can be seen when choosing large windows for clumping, where many of the strong association signals are removed due to long distance LD induced by the population structure. However, even when using a smaller maximum distance, the signals that remained after clumping were weaker compared to using standard LD clumping.

## Code and data availability

We implemented pruning and clumping based on adjusted LD in the software PCAone [52], which can be downloaded at https://github.com/Zilong-Li/PCAone. Example usage of pruning from PLINK files is

~~~
PCAone -b plink -k 3 -n 20 --ld-r2 0.8 --ld-window 1000000 --maf 0.05
~~~

where binary PLINK files with prefix `plink`, -k 3 principal components are used, -n 20 CPU threads, and with the PCA perform in memory based on sites with a minor allele frequency above 5% (–maf 0.05). For pruning, a *r*^2^ threshold of 0.8 is used within a window size of 1Mb.

Example usage of clumping is

~~~
PCAone -B pcaone.residuals --clump plink.assoc
  --clump-p1 5e-8 --clump-p2 1e-6
  --clump-r2 0.01 --clump-bp 10000000
~~~

where `pcaone.residuals` is a binary file outputted by PCAone in the pruning step, and `plink.assoc` text file stores the association testing result, and the rest options define thresholds used for clumping.

## Notes

### Competing Interest Statement

The authors have declared no competing interest.

