## Supplementary material for "Measuring linkage disequilibrium and improvement of pruning and clumping in structured populations"

### S1 Appendix

*Proof of Theorem 1.* Let $Z_s$ and $Z_t$ be random variables in $\{1, \cdots, k\}$ that denote the ancestral origin of a haplotype of the SNPs $s$ and $t$, respectively, of a random individual. Then,

$$\begin{aligned}(D_{\text{adj}})_{st} &= \frac{1}{2}\mathbb{E}[\text{Cov}(G_s, G_t | Z_s, Z_t)] \\ &= \frac{1}{2}\sum_{\ell_s=1}^{k}\sum_{\ell_t=1}^{k}\mathbb{P}(Z_s = \ell_s, Z_t = \ell_t)\text{Cov}(G_s, G_t | Z_s = \ell_s, Z_t = \ell_t).\end{aligned}$$

As $\text{Cov}(G_s, G_t | Z_s = \ell_s, Z_t = \ell_t) = 0$ when $\ell_s \neq \ell_t$,

$$\begin{aligned}&\frac{1}{2}\sum_{\ell_s=1}^{k}\sum_{\ell_t=1}^{k}\mathbb{P}(Z_s = \ell_s, Z_t = \ell_t)\text{Cov}(G_s, G_t | Z_s = \ell_s, Z_t = \ell_t) \\ &= \frac{1}{2}\sum_{\ell=1}^{k}\mathbb{P}(Z_s = \ell, Z_t = \ell)\text{Cov}(G_s, G_t | Z_s = \ell, Z_t = \ell).\end{aligned}$$

Letting $w_{st}^{\ell} = \mathbb{P}(Z_s = \ell, Z_t = \ell)$, then the first result follows and $\sum_{\ell=1}^{k} w_{st}^{\ell} \leq 1$. Equality holds if and only if $\mathbb{P}(Z_s = \ell, Z_t = \ell) = \mathbb{P}(Z_s = \ell)$, which is the case if the individual has all genetic material from one ancestral population. □

**Lemma S1.** *Let $D'$ be the matrix with entries*

$$D'_{st} = \frac{1}{2(n-k)}\sum_{i=1}^{n}(R'_{is} - \overline{R'}_{\cdot s})(R'_{it} - \overline{R'}_{\cdot t}),$$

*where $R' = (I - P')G$ and $P'$ is an arbitrary projection matrix. Then, $D'$ in matrix form is given by*

$$D' = \frac{1}{2(n-k)}G^{\mathrm{T}}(I - P')(I - J/n)(I - P')G.$$

*Moreover, if $P'\mathbf{1} = \mathbf{1}$, where $\mathbf{1}$ is the column vector of ones, then*

$$D' = \frac{1}{2(n-k)}G^{\mathrm{T}}(I - P')G.$$

*Proof.* From the definition of $D'$, we have that

$$\begin{aligned}
D'_{st} &= \frac{1}{2(n-k)}\sum_{i=1}^{n}[R'_{is}-\overline{R'_{\cdot s}}][R'_{it}-\overline{R'_{\cdot t}}]\\
&= \frac{1}{2(n-k)}\sum_{i=1}^{n}[(I-P')_{i\cdot}G_{\cdot s}-\overline{(I-P')_{i\cdot}G_{\cdot s}}][(I-P')_{i\cdot}G_{\cdot t}-\overline{(I-P')_{i\cdot}G_{\cdot t}}]\\
&= \frac{1}{2(n-k)}\sum_{i=1}^{n}[(I-P')_{i\cdot}G_{\cdot s}-\overline{(I-P')}G_{\cdot s}][(I-P')_{i\cdot}G_{\cdot t}-\overline{(I-P')}G_{\cdot t}]\\
&= \frac{1}{2(n-k)}\sum_{i=1}^{n}[((I-P')_{i\cdot}-\overline{(I-P')})G_{\cdot s}][((I-P')_{i\cdot}-\overline{(I-P')})G_{\cdot t}]\\
&= \frac{1}{2(n-k)}\sum_{i=1}^{n}(I-P')_{i\cdot}G_{\cdot s}(I-P')_{i\cdot}G_{\cdot t}-\frac{n}{2(n-k)}\overline{(I-P')}G_{\cdot s}\overline{(I-P')}G_{\cdot t}\,,
\end{aligned}$$

where $\overline{(I-P')}=\sum_{i=1}^{n}(I-P')_{i\cdot}/n=\mathbf{1}^{\mathrm{T}}(I-P')/n$. We consider these terms individually. As $P'$ is a projection matrix, it is symmetric and idempotent. Hence, $(I-P')$ is also symmetric and $(I-P')^2=I-P'$. Then, by using these properties,

$$\begin{aligned}
\frac{1}{2(n-k)}\sum_{i=1}^{n}(I-P')_{i\cdot}G_{\cdot s}(I-P')_{i\cdot}G_{\cdot t} &= \frac{1}{2(n-k)}\sum_{i=1}^{n}G_{\cdot s}{}^{\mathrm{T}}(I-P')_{i\cdot}{}^{\mathrm{T}}(I-P')_{i\cdot}G_{\cdot t}\\
&= \frac{1}{2(n-k)}G_{\cdot s}{}^{\mathrm{T}}(I-P')G_{\cdot t}\\
&= \frac{1}{2(n-k)}G_{\cdot s}{}^{\mathrm{T}}(I-P')(I-P')G_{\cdot t}
\end{aligned}$$

and, by expanding $\overline{(I-P')}$,

$$\begin{aligned}
\frac{n}{2(n-k)}\overline{(I-P')}G_{\cdot s}\overline{(I-P')}G_{\cdot t} &= \frac{1}{2n(n-k)}\mathbf{1}^{\mathrm{T}}(I-P')G_{\cdot s}\mathbf{1}^{\mathrm{T}}(I-P')G_{\cdot t}\\
&= \frac{1}{2n(n-k)}G_{\cdot s}{}^{\mathrm{T}}(I-P')\mathbf{1}\mathbf{1}^{\mathrm{T}}(I-P')G_{\cdot t}\\
&= \frac{1}{2n(n-k)}G_{\cdot s}{}^{\mathrm{T}}(I-P')J(I-P')G_{\cdot t}\,.
\end{aligned}$$

Therefore, $D'$ can be written as

$$D'=\frac{1}{2(n-k)}G^{\mathrm{T}}(I-P')(I-J/n)(I-P')G.$$

For the case of interest where $P'\mathbf{1}=\mathbf{1P'}=\mathbf{1}$, $\overline{(I-P')}=0$ and then

$$D'=\frac{1}{2(n-k)}G^{\mathrm{T}}(I-P')G.$$

□

To study the asymptotic behaviour of the ancestry adjusted LD measure when the number of individuals and/or SNPs is large, we impose some conditions: 1) Large number of individuals, $n\to\infty$. Individuals randomly choose their admixture proportions from a common pool representing the admixture population as such. Specifically, we assume the row vectors of $Q$, representing the admixture proportions of the different individuals, are independent draws from a common distribution. For example, if the population is homogeneous, then $k = 1$ and each row of $Q$ is just the number one. Hence, the common distribution is the degenerate distribution that always returns one. If the population consists of two homogeneous sub-populations, then $k = 2$ and each row of $Q$ is either $(1, 0)$ or $(0, 1)$, indicating whether the ancestry of an individual comes from the first or second sub-population, and the common distribution specifies the probabilities by which these two vectors are drawn. 2) Large number of SNPs, $m \to \infty$. To validate the PCA approach, we make the simplifying assumption that SNP frequencies for different SNPs (different columns in $F$) are independent and identically distributed [1]. In particular, this assumption implies that $\widehat{P}^{n,m} \to P$ (an $n \times n$ matrix) pointwise as $m \to \infty$ with $n$ fixed. Independence is a reasonable assumption as the vast majority of SNP pairs are not in LD. Furthermore, as LD tends to be local and not stretch far, the convergence result quoted above would also be valid with some LD [1].

*Proof of Theorem 2.* From (8) in the main paper it follows that

$$(\widehat{D}^{n,m}_{\text{adj}})_{st} \to \frac{1}{2(n-k)} G^{\mathrm{T}}_{\cdot s}(I-P)(I-J/n)(I-P)G_{\cdot t}.$$

Since the vector $\mathbf{1}$ is in the column space of $Q$ by assumption, then $P\mathbf{1} = \mathbf{1}$, and consequently $PJ = J$ and $JP = J$. It follows that $(I-P)(I-J/n)(I-P) = I-P$, and the first convergence result follows.

For the second part, note that $\widehat{D}^{m,n}_{\text{adj}}$ is bounded,

$$\begin{aligned}
|(\widehat{D}^{n,m}_{\text{adj}})_{st}|^2 &= \left| \frac{1}{2(n-k)} G^{\mathrm{T}}_{\cdot s}(I-\widehat{P}^{n,m})(I-J/n)(I-\widehat{P}^{n,m})G_{\cdot t} \right|^2 \\
&= \left\| \frac{1}{2(n-k)} G^{\mathrm{T}}_{\cdot s}(I-\widehat{P}^{n,m})(I-J/n)(I-\widehat{P}^{n,m})G_{\cdot t} \right\|^2_{\mathcal{F}} \\
&\le \frac{1}{4(n-k)^2} \|G_{\cdot s}\|^2_{\mathcal{F}} \|I-\widehat{P}^{n,m}\|^2_{\mathcal{F}} \|I-J/n\|^2_{\mathcal{F}} \|I-\widehat{P}^{n,m}\|^2_{\mathcal{F}} \|G_{\cdot t}\|^2_{\mathcal{F}} \\
&\le \frac{1}{4(n-k)^2} 4n(n-k) \frac{(n-1)^2+n-1}{n} (n-k)4n \\
&= 4n\left((n-1)^2+n-1\right).
\end{aligned}$$

Then, as $\widehat{D}^{n,m}_{\text{adj}}$ is uniformly bounded in $m$, by the dominated convergence theorem, $\mathbb{E}\left[(\widehat{D}^{n,m}_{\text{adj}})_{st}\right] \to \mathbb{E}\left[(\widehat{D}_{\text{adj}})_{st}\right]$ as $m \to \infty$.

It remains to calculate the expectation of the limit. We have

$$\begin{aligned}
\mathbb{E}[(\widehat{D}_{\text{adj}})_{st}] &= \mathbb{E}\left[\frac{1}{2(n-k)}\sum_{i=1}^{n}\left((I-P)_{i\cdot}-\overline{(I-P)}\right)G_{\cdot s}\left((I-P)_{i\cdot}-\overline{(I-P)}\right)G_{\cdot t}\right] \\
&= \frac{1}{2(n-k)}\sum_{i=1}^{n}\left((I-P)_{i\cdot}-\overline{(I-P)}\right)\mathbb{E}[G_{\cdot s}G_{\cdot t}{}^{\mathrm{T}}]\left((I-P)_{i\cdot}-\overline{(I-P)}\right)^{\mathrm{T}} \\
&= \frac{1}{2(n-k)}\sum_{i=1}^{n}(I-P)_{i\cdot}\mathbb{E}[G_{\cdot s}G_{\cdot t}{}^{\mathrm{T}}](I-P)_{i\cdot}{}^{\mathrm{T}},
\end{aligned}$$

where we used that the vector $\mathbf{1}$ is in the column space of $Q$, hence $P\mathbf{1} = \mathbf{1}$ and $\overline{(I-P)} = 0$. As $\mathbb{E}[G_{\cdot s}] = 2QF_{\cdot s}$ and $(I-P)Q = 0$, we get $(I-P)\mathbb{E}[G_{\cdot s}] = 0$. Hence, $\mathbb{E}[G_{\cdot s}G_{\cdot t}{}^{\mathrm{T}}]$ is equivalent to the $n \times n$ matrix given by $H_{st} := \mathrm{Cov}[G_{\cdot s}G_{\cdot t}{}^{\mathrm{T}}]$.

For the case where $G_{is}$ is independent of $G_{it}$ for every individual, $(H_{st})_{ii} = 0$ for $i = 1, \ldots, n$ and then $\mathbb{E}[(\widehat{D}_{\text{adj}})_{st}] = 0$. □

**Lemma S2.** *Let $Q$ be is given as in* (10)*. Then, $C_{st}$ is the pooled covariance of the sub-populations given by*

$$C_{st} = \frac{\sum_{\ell=1}^{k}(n_\ell - 1)(p_{st}^{\ell} - p_s^{\ell}p_t^{\ell})}{2(n-k)},$$

*where $n_\ell$ is the number of individuals from sub-population $\ell$.*

*Proof.* As $Q$ is given as in (10), we can compute $Q(Q^{\mathrm{T}}Q)^{-1}Q^{\mathrm{T}}$ to see that the projection matrix $P$ is given by

$$P = \begin{pmatrix} P_1 & 0 & \ldots & 0 \\ 0 & P_2 & \ldots & 0 \\ \vdots & \vdots & \ddots & \vdots \\ 0 & 0 & \ldots & P_k \end{pmatrix}, \tag{S1}$$

where $P_\ell$ is an $n_\ell \times n_\ell$ matrix with $1/n_\ell$ on each element. Hence, $\overline{I-P} = 0$ and we can write $C_{st}$ as

$$\begin{aligned}
C_{st} &= \frac{1}{2(n-k)}\sum_{i=1}^{n}((I-P)_{i\cdot})H_{st}((I-P)_{i\cdot})^{\mathrm{T}} \\
&= \frac{1}{2(n-k)}\mathrm{TR}((I-P)H_{st}(I-P)) = \frac{1}{2(n-k)}(\mathrm{TR}(H_{st}) - \mathrm{TR}(PH_{st})) \\
&= \frac{1}{2(n-k)}\left(\sum_{i=1}^{n}\mathrm{Cov}(G_{is},G_{it}) - \sum_{i=1}^{n}\sum_{\ell=1}^{k}\mathrm{Cov}(G_{is},G_{it})\frac{Q_{i\ell}}{n_\ell}\right).
\end{aligned}$$

Then, as $\mathrm{Cov}(G_{is}, G_{it}) = p_{st}^{\ell} - p_s^{\ell}p_t^{\ell}$ if the individual $i$ belongs to the sub-population $\ell$, we have that

$$\frac{1}{2(n-k)}\left(\sum_{i=1}^{n}\mathrm{Cov}(G_{is},G_{it})-\sum_{i=1}^{n}\sum_{\ell=1}^{k}\mathrm{Cov}(G_{is},G_{it})\frac{Q_{i\ell}}{n_\ell}\right)$$
$$=\frac{1}{2(n-k)}\sum_{\ell=1}^{k}(n_\ell-1)(p^\ell_{st}-p^\ell_s p^\ell_t).$$

□

*Proof of Theorem 3.* With $Q$ known, then $P=Q(Q^{\mathrm{T}}Q)^{-1}Q^{\mathrm{T}}$ and $P$ needs not be estimated. We consider $n\to\infty$ and note that this implies the dimension of the matrices, in particular $P$, increases with $n$. However, we suppress the dimensions in the notation for convenience. By Lemma S1, we can decompose $\widehat{D}_{\mathrm{adj}}$ as

$$(\widehat{D}_{\mathrm{adj}})_{st}=\frac{1}{2}\left(\frac{1}{n-k}{G_{\cdot s}}^{\mathrm{T}}(I-P)G_{\cdot t}-\frac{n}{n-k}\overline{(I-P)G_{\cdot s}}\,\overline{(I-P)G_{\cdot t}}\right). \quad \text{(S2)}$$

Consider first $\frac{1}{n-k}{G_{\cdot s}}^{\mathrm{T}}(I-P)G_{\cdot t}$. As

$$\frac{1}{n-k}{G_{\cdot s}}^{\mathrm{T}}(I-P)G_{\cdot t}=\frac{1}{n-k}{G_{\cdot s}}^{\mathrm{T}}G_{\cdot t}-\frac{1}{n-k}{G_{\cdot s}}^{\mathrm{T}}PG_{\cdot t},$$

we look at each term individually. The fact that the rows of $G$ are i.i.d., ensures we can use the Law of Large Numbers. First, consider $\frac{1}{n-k}{G_{\cdot s}}^{\mathrm{T}}G_{\cdot t}$:

$$\frac{1}{n-k}{G_{\cdot s}}^{\mathrm{T}}G_{\cdot t}=\frac{1}{n-k}\sum_{i=1}^{n}G_{is}G_{it}\quad\to\quad\mathbb{E}[G_{1s}G_{1t}] \quad \text{(S3)}$$

as $n\to\infty$ (that is, for almost all realisations of $Q$ and $G$, convergence holds). Secondly, we decompose $\frac{1}{n-k}G^{\mathrm{T}}PG$ using that $P=Q(Q^{\mathrm{T}}Q)^{-1}Q^{\mathrm{T}}$. Therefore,

$$\frac{1}{n-k}{G_{\cdot s}}^{\mathrm{T}}PG_{\cdot t}=\frac{1}{n-k}{G_{\cdot s}}^{\mathrm{T}}Q(Q^{\mathrm{T}}Q)^{-1}Q^{\mathrm{T}}G_{\cdot t}$$
$$=\frac{1}{n-k}{G_{\cdot s}}^{\mathrm{T}}Q\left(\frac{1}{n-k}Q^{\mathrm{T}}Q\right)^{-1}\frac{1}{n-k}Q^{\mathrm{T}}G_{\cdot t}\,.$$

We investigate $\frac{1}{n-k}G^{\mathrm{T}}Q$ and $(\frac{1}{n-k}Q^{\mathrm{T}}Q)^{-1}$ individually, noticing that $\frac{1}{n-k}Q^{\mathrm{T}}G$ is just the transpose of $\frac{1}{n-k}G^{\mathrm{T}}Q$. Starting with the $k$-dimensional vector $\frac{1}{n-k}{G_{\cdot s}}^{\mathrm{T}}Q$ and using the Law of Large Numbers,

$$\left(\frac{1}{n-k}{G_{\cdot s}}^{\mathrm{T}}Q\right)_\ell=\frac{1}{n-k}{G_{\cdot s}}^{\mathrm{T}}Q_{\cdot\ell}=\frac{1}{n-k}\sum_{i=1}^{n}G_{is}Q_{i\ell}$$
$$\to\mathbb{E}[G_{1s}Q_{1\ell}]=2\mathbb{E}[Q_{1\cdot}F_{\cdot s}Q_{1\ell}]$$

as $n\to\infty$. Then, for $n\to\infty$,

$$\frac{1}{n-k}{G_{\cdot s}}^{\mathrm{T}}Q\to 2\mathbb{E}[Q_{1\cdot}F_{\cdot s}Q_{1\cdot}]. \quad \text{(S4)}$$

For the $k \times k$ matrix $\left(\frac{1}{n-k} Q^{\mathrm{T}} Q\right)^{-1}$, we have for $n \to \infty$,

$$\left(\frac{1}{n-k} Q^{\mathrm{T}} Q\right)^{-1} \to \mathbb{E}[Q_{1\cdot}{}^{\mathrm{T}} Q_{1\cdot}]^{-1}. \tag{S5}$$

Hence, returning to $\frac{1}{n-k} G^{\mathrm{T}} P G$ and using equations (S4) and (S5),

$$\begin{aligned} \frac{1}{n-k} G_{\cdot s}{}^{\mathrm{T}} P G_{\cdot t} &\to 4\mathbb{E}[Q_{1\cdot} F_{\cdot s} Q_{1\cdot}](\mathbb{E}[Q_{1\cdot}{}^{\mathrm{T}} Q_{1\cdot}])^{-1} \mathbb{E}[Q_{1\cdot} F_{\cdot t} Q_{1\cdot}]^{\mathrm{T}} \\ &= 4\mathbb{E}[Q_{1\cdot} F_{\cdot s} Q_{1\cdot} F_{\cdot t}] \end{aligned} \tag{S6}$$

for $n \to \infty$. By combining equation (S3) and (S6), we get that

$$\frac{1}{n-k} G_{\cdot s}{}^{\mathrm{T}} G_{\cdot t} - \frac{1}{n-k} G_{\cdot s}{}^{\mathrm{T}} P G_{\cdot t} \to \mathbb{E}[G_{1s} G_{1t}] - 4\mathbb{E}[Q_{1\cdot} F_{\cdot s} Q_{1\cdot} F_{\cdot t}] \tag{S7}$$

for $n \to \infty$.

For the second term,

$$\begin{aligned} \overline{(I-P)G_{\cdot s}}\,\overline{(I-P)G_{\cdot t}} &= \frac{1}{n}\sum_{i=1}^{n}(I-P)_{i\cdot} G_{\cdot s} \frac{1}{n}\sum_{j=1}^{n}(I-P)_{j\cdot} G_{\cdot t} \\ &= \left[\left(\frac{1}{n}\mathbf{1}^{\mathrm{T}} - \frac{1}{n}\sum_{i=1}^{n} P_{i\cdot}\right) G_{\cdot s}\right]\left[\left(\frac{1}{n}\mathbf{1}^{\mathrm{T}} - \frac{1}{n}\sum_{j=1}^{n} P_{j\cdot}\right) G_{\cdot t}\right]. \end{aligned} \tag{S8}$$

We calculate individually $\frac{1}{n}\mathbf{1}^{\mathrm{T}} G_{\cdot s}$ and $\frac{1}{n}\sum_{i=1}^{n} P_{i\cdot} G_{\cdot s}$, and use the Law of Large Numbers over the rows of $G$. Consider $\frac{1}{n}\mathbf{1}^{\mathrm{T}} G_{\cdot s}$:

$$\frac{1}{n}\mathbf{1}^{\mathrm{T}} G_{\cdot s} = \frac{1}{n}\sum_{i=1}^{n} G_{is} \to \mathbb{E}[G_{1s}] = 2\mathbb{E}[Q_{1\cdot} F_{\cdot s}] \tag{S9}$$

for $n \to \infty$. For $\frac{1}{n}\sum_{i=1}^{n} P_{i\cdot} G_{\cdot s}$, using the same techniques as in the first term and by decomposing $P$,

$$\begin{aligned} \frac{1}{n}\sum_{i=1}^{n} P_{i\cdot} G_{\cdot s} &= \frac{1}{n}\sum_{i=1}^{n} Q_{i\cdot}\left(\frac{1}{n} Q^{\mathrm{T}} Q\right)^{-1} \frac{1}{n} Q^{\mathrm{T}} G_{\cdot s} \\ &\to \mathbb{E}[Q_{1\cdot}](\mathbb{E}[Q_{1\cdot}{}^{\mathrm{T}} Q_{1\cdot}])^{-1} 2\mathbb{E}[Q_{1\cdot}{}^{\mathrm{T}} Q_{1\cdot} F_{\cdot s}] \\ &= 2\mathbb{E}[Q_{1\cdot} F_{\cdot s}] \end{aligned} \tag{S10}$$

for $n \to \infty$. Hence, combining equations (S9) and (S10),

$$\left(\frac{1}{n}\mathbf{1}^{\mathrm{T}} - \frac{1}{n}\sum_{j=1}^{n} P_{j\cdot}\right) G_{\cdot s} \to 0$$

as $n \to \infty$. Noticing that the same calculations can be done for $G_{\cdot t}$, we use equation (S8) to conclude that

$$\overline{(I-P)G_{\cdot s}}\,\overline{(I-P)G_{\cdot t}} \to 0$$

as $n \to \infty$.

Therefore,

$$
\begin{aligned}
(\widehat{D}_{\mathrm{adj}})_{st} &= \frac{1}{2}\left(\frac{1}{n-k}G_{\cdot s}{}^{\mathrm{T}}(I-P)G_{\cdot t} - \frac{n}{n-k}\overline{(I-P)}G_{\cdot s}\overline{(I-P)}G_{\cdot t}\right) \\
&\to \frac{1}{2}\left(\mathbb{E}[G_{1s}G_{1t}] - 4\mathbb{E}[Q_{1\cdot}F_{\cdot s}Q_{1\cdot}F_{\cdot t}]\right) \\
&= \frac{1}{2}\left(\mathrm{Cov}(G_{1s}, G_{1t}) - \mathrm{Cov}(\mathbb{E}[G_{1s}|\Pi], \mathbb{E}[G_{1t}|\Pi])\right) \\
&= \frac{1}{2}\mathbb{E}[\mathrm{Cov}(G_{1s}, G_{1t}|\Pi)] = (D_{\mathrm{adj}})_{st} \qquad \text{(S11)}
\end{aligned}
$$

as $n \to \infty$, where in the second equality after the limit we use the Law of total covariance. □

### S2 Supplementary figures

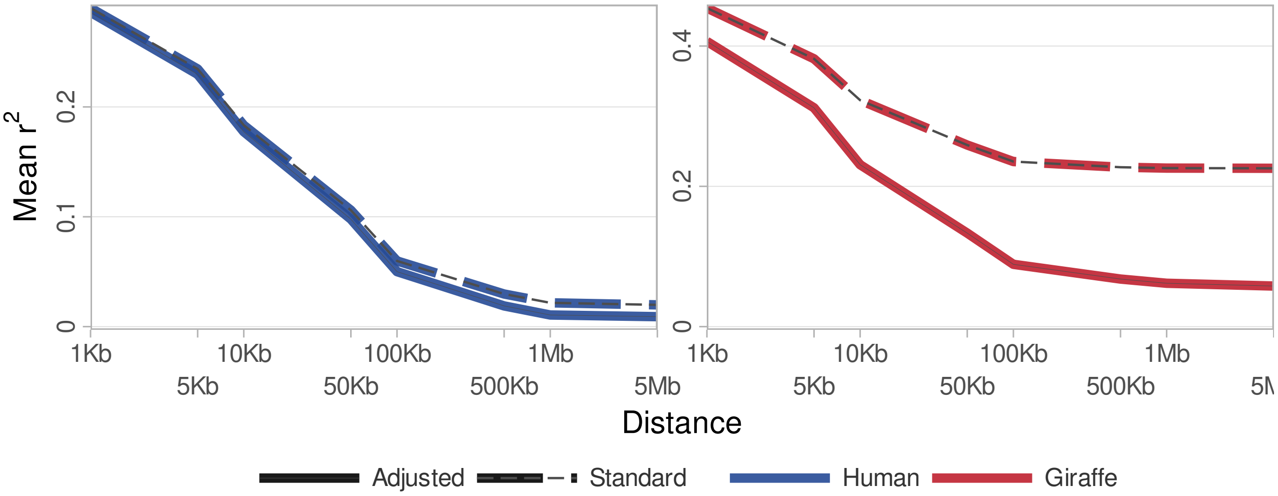


Figure S1: LD decay curves of standard $r^2$ for the data sets jointly, with no sample size correction

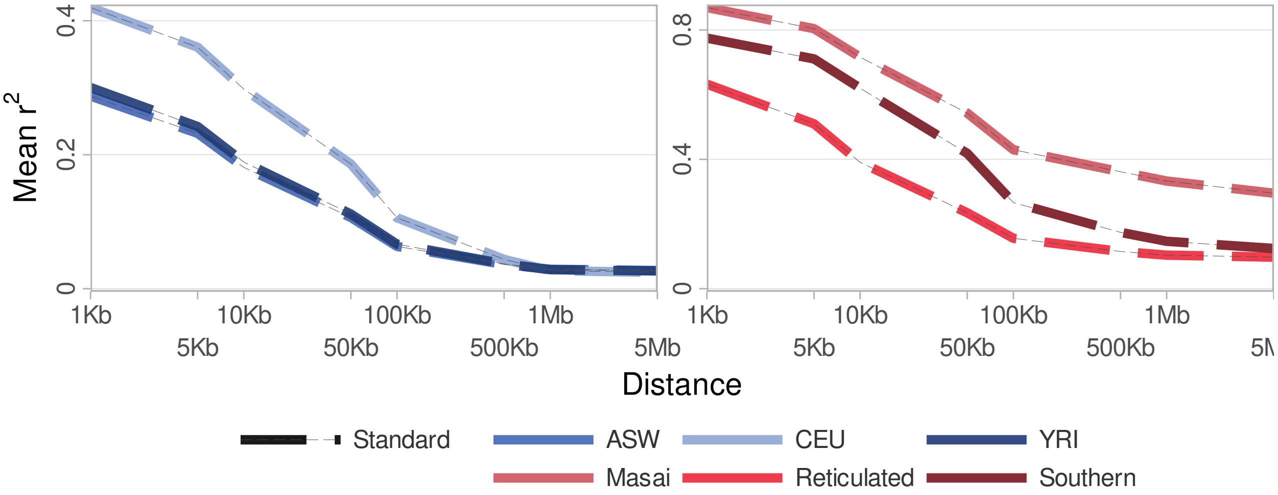


Figure S2: LD decay curves of standard $r^2$ for each of the constituent populations considered separately.

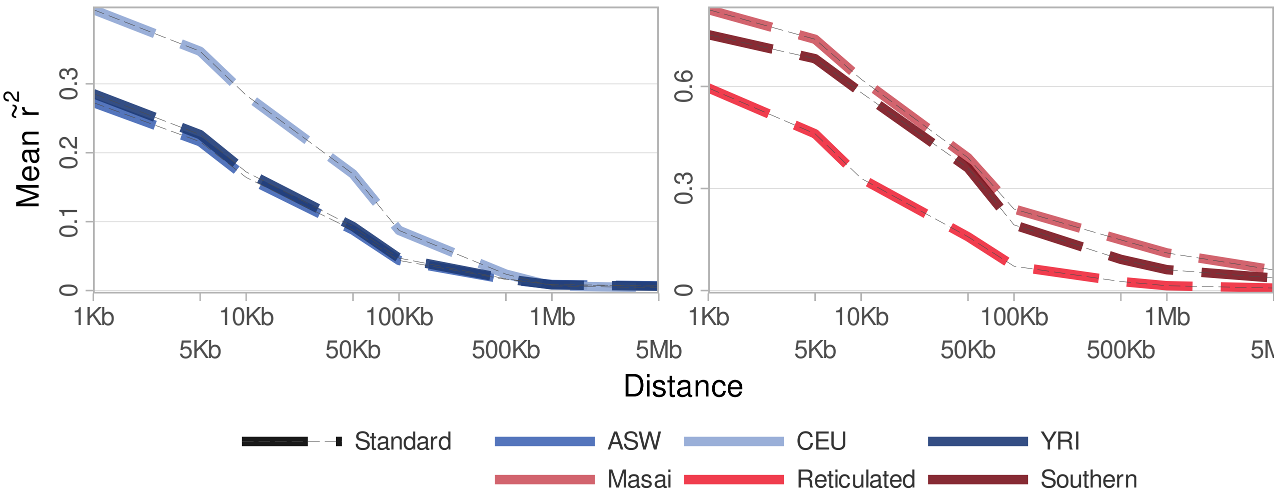


Figure S3: LD decay curves of standard $\tilde{r}^2$ for each of the constituent populations considered separately.

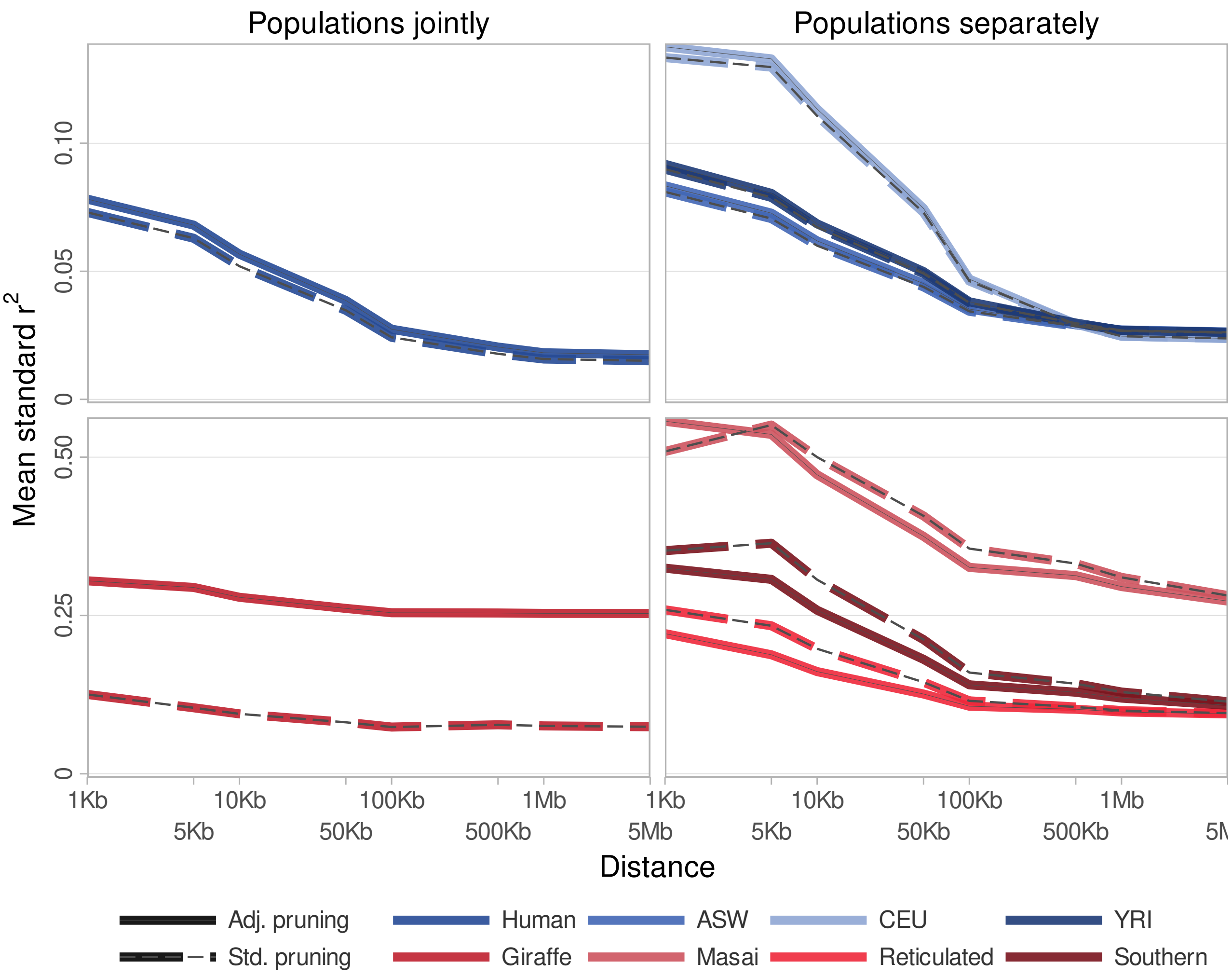


Figure S4: Standard LD decay curves after pruning based on either standard or adjusted $r^2$. The joint datasets are shown, as well as LD decay curves for the constituent populations extracted from the pruned datasets.

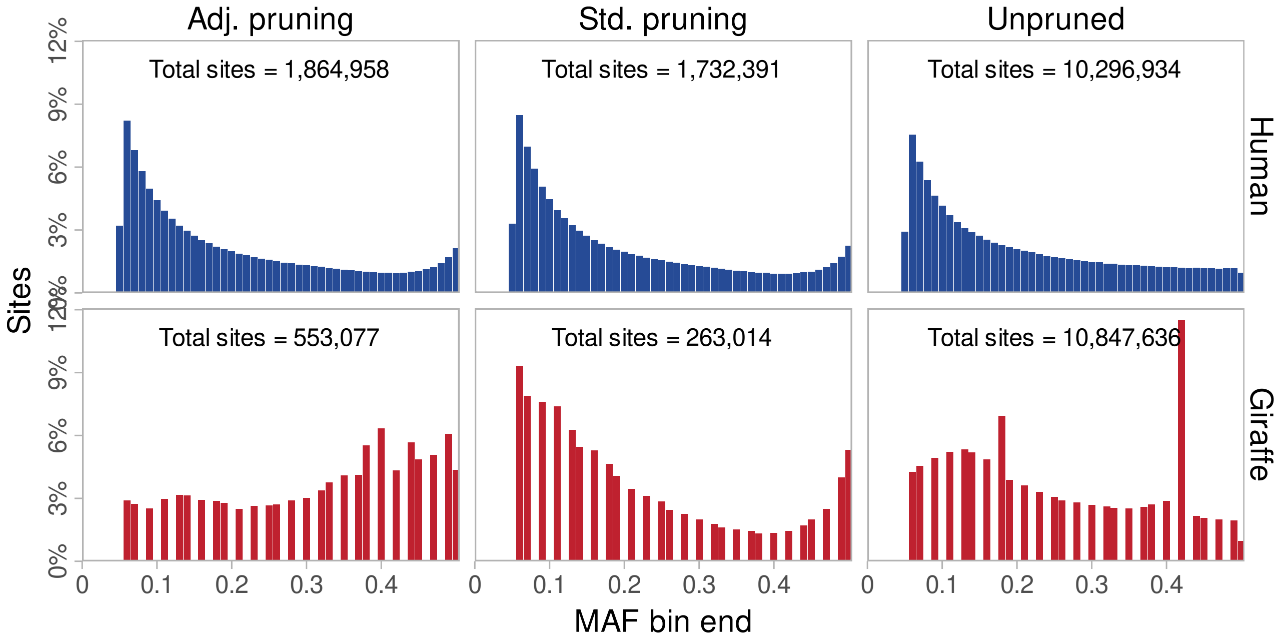


Figure S5: Distribution of minor allele frequencies before and after pruning based on either standard or adjusted $r^2$. Total number of sites in the data overlaid.

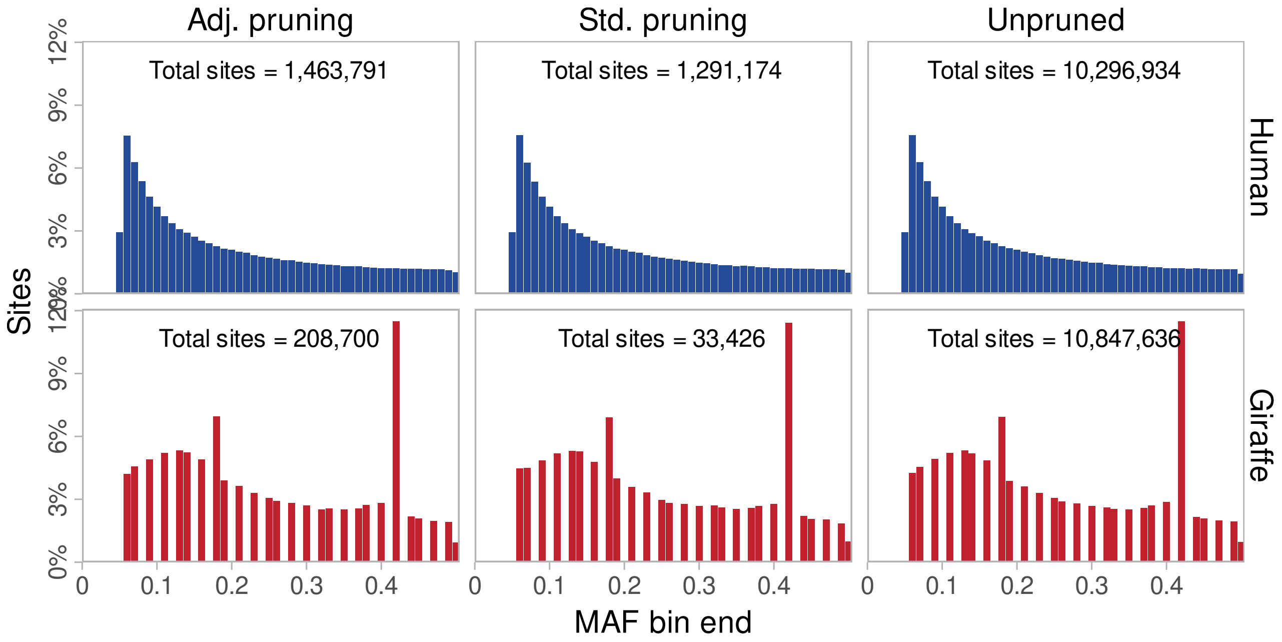


Figure S6: Distribution of minor allele frequencies before and after pruning based on either standard or adjusted $r^2$, having resampled the pruned data to get a MAF distribution approximating the unpruned distribution. Total number of sites in the data overlaid.

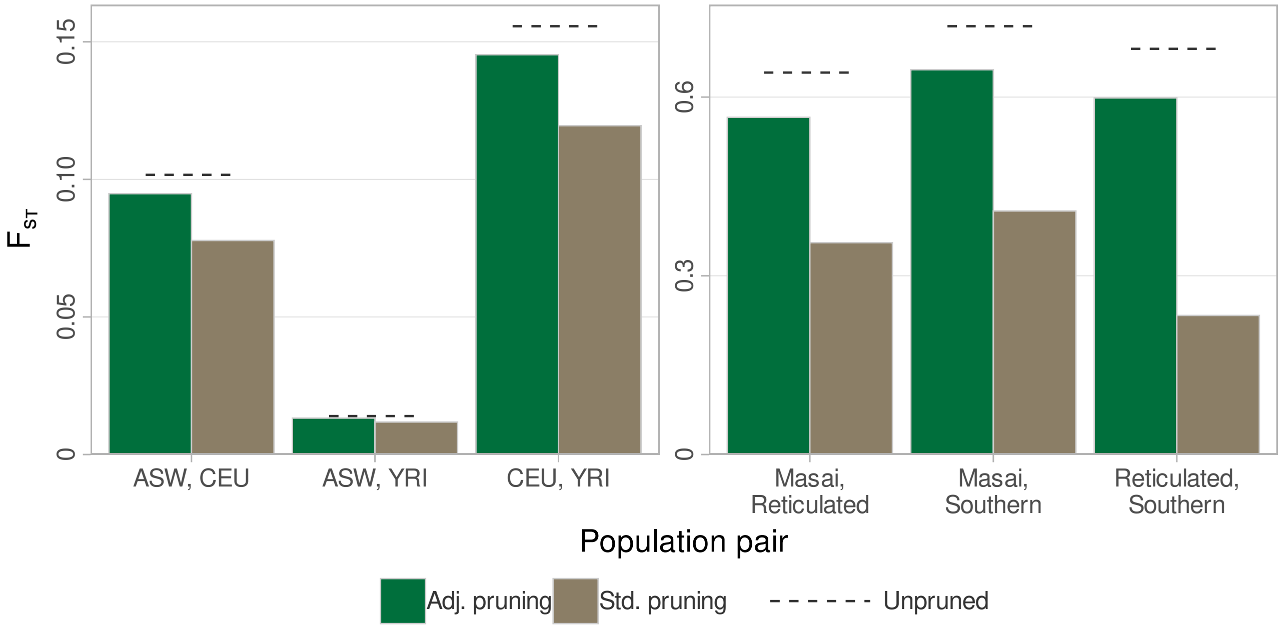


Figure S7: $F_{ST}$ after pruning and resampling to match the unpruned MAF distribution, i.e. based on the data shown in fig. S6. $F_{ST}$ shown for each pair of populations, with the unpruned $F_{ST}$ shown for comparison.

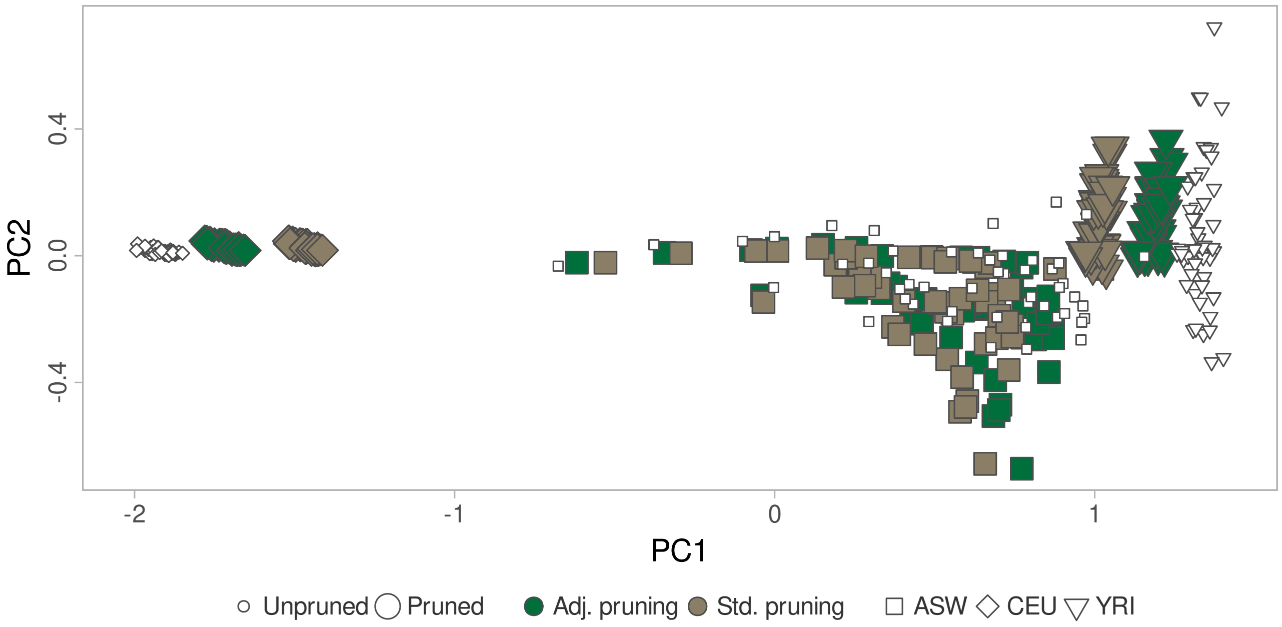


Figure S8: PCA after pruning on the human dataset, with the unpruned data included for comparison. Eigenvectors scaled by corresponding eigenvalue shown.

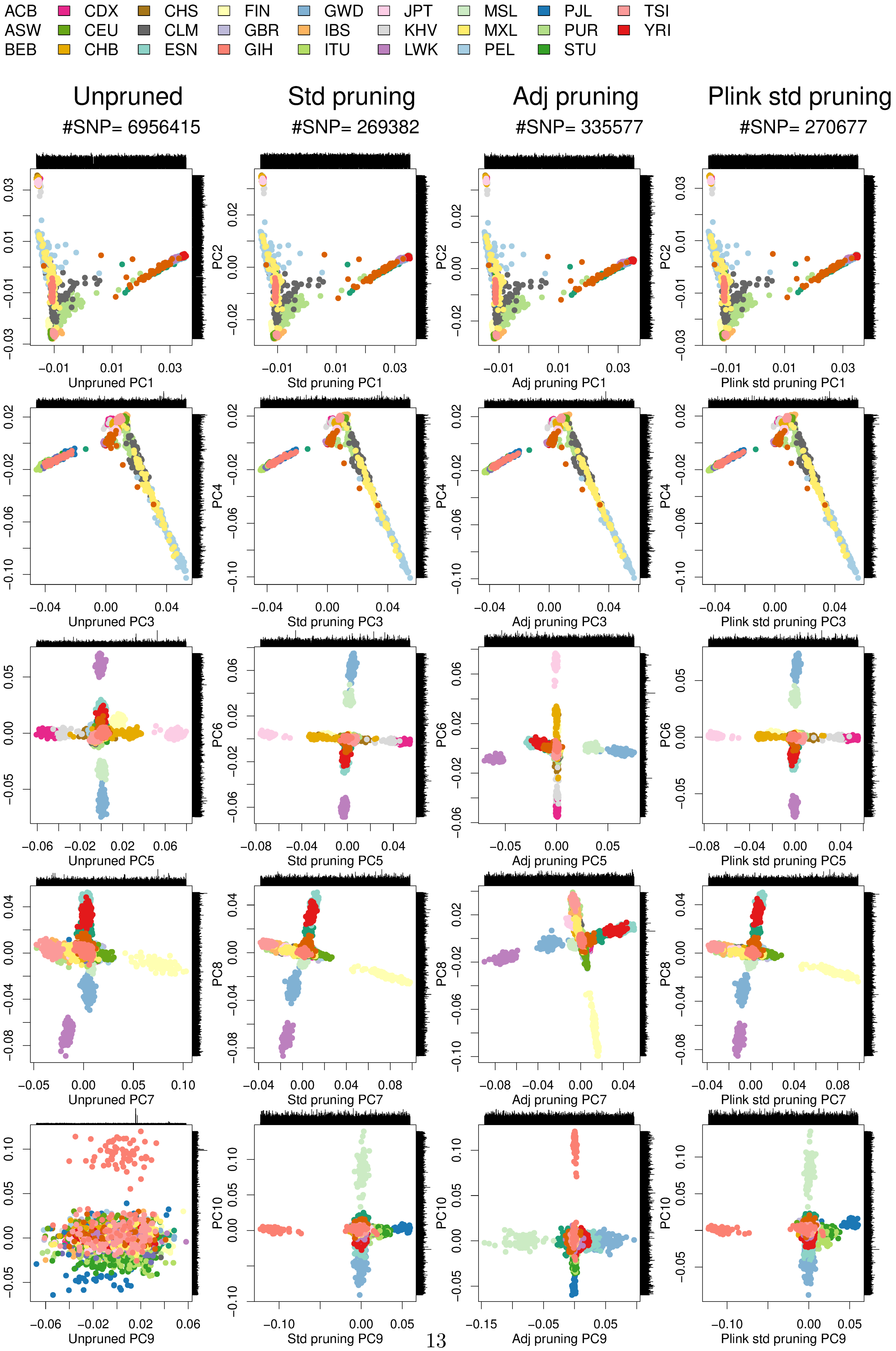


Figure S9: Top 1-10 PCs of 1000 Genomes data using different pruning schemes. The SNP loadings are shown on the side of the plot.

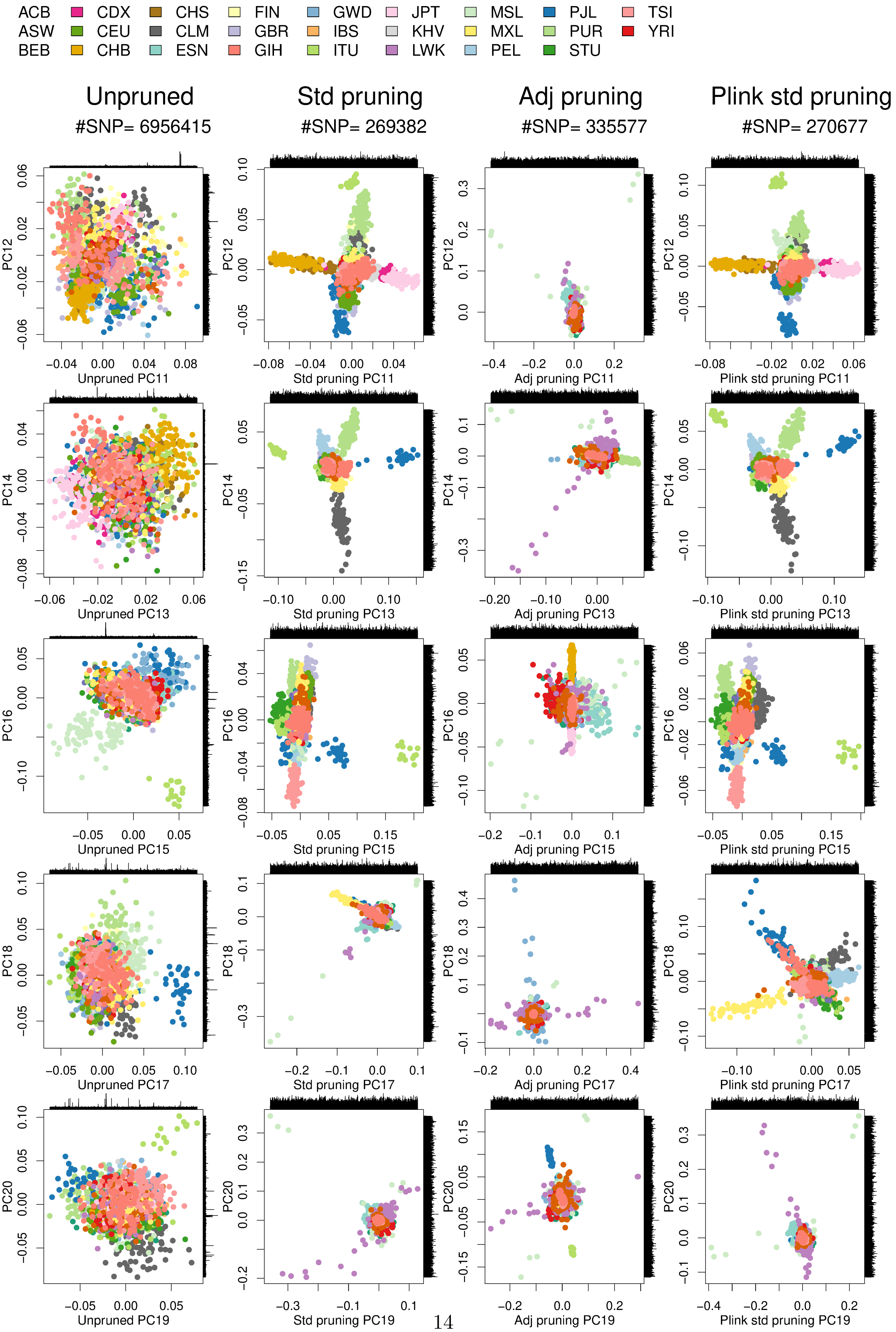




Figure S10: Top 11-20 PCs of 1000 Genomes data using different pruning schemes. The SNP loadings are shown on the side of the plot.

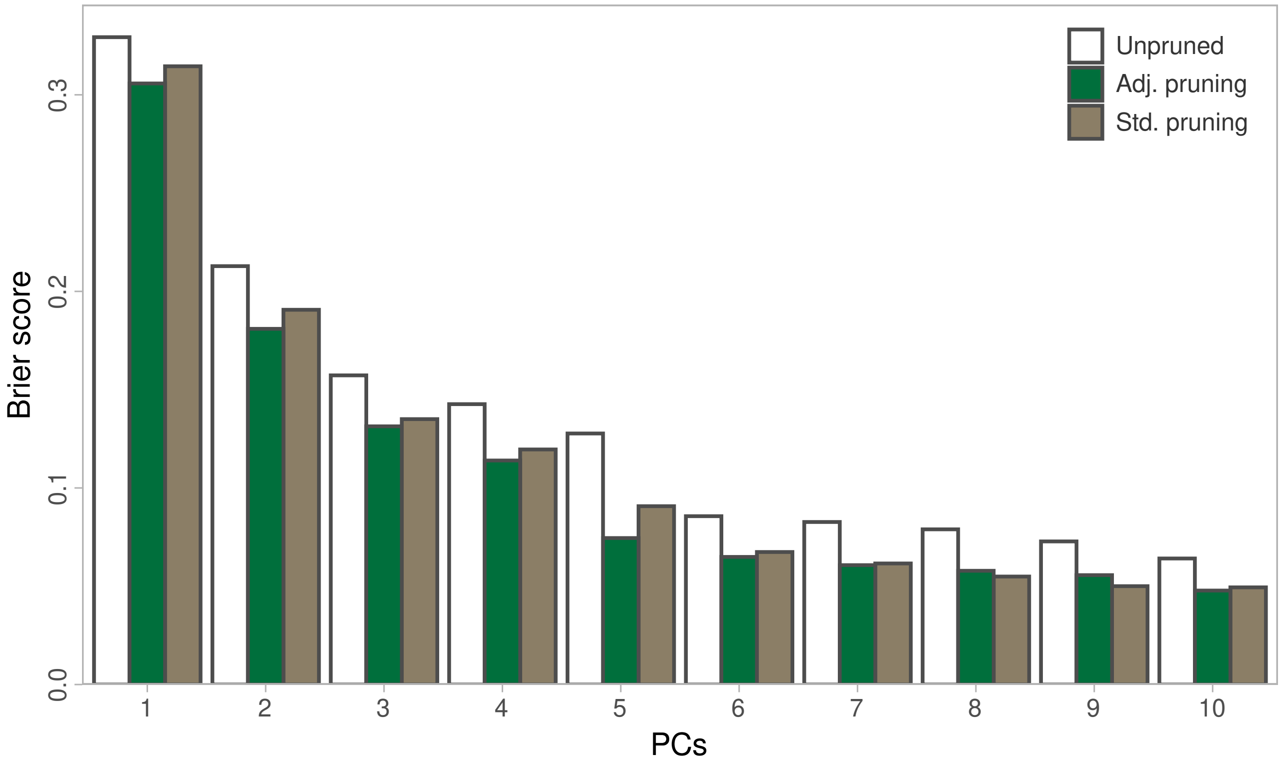


Figure S11: Brier scores from `mclust` using various numbers of PCs on the 1000G data, comparing pruning methods.

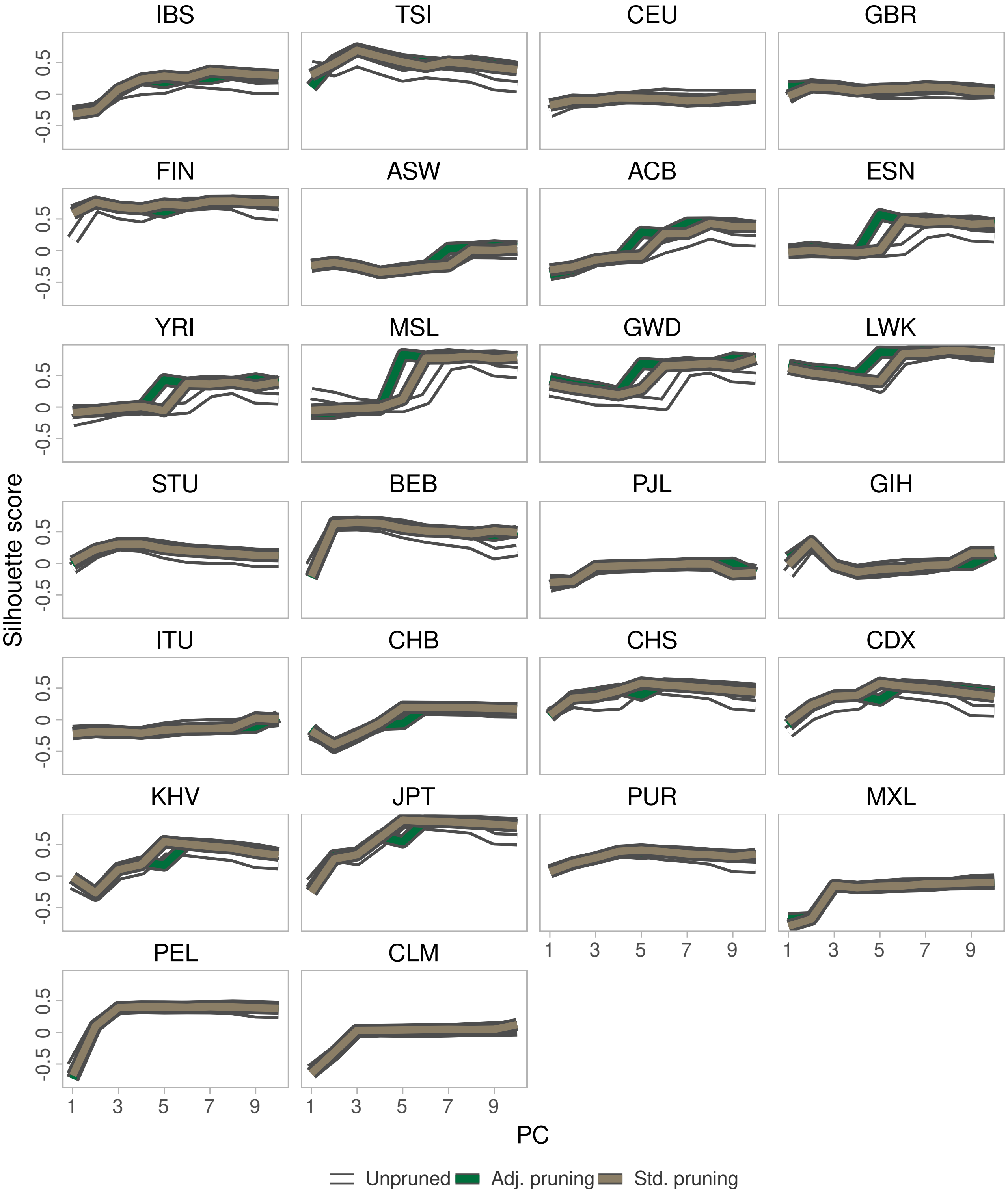


Figure S12: Silhouette scores averaged by population using various numbers of PCs on the 1000G data, comparing pruning methods.

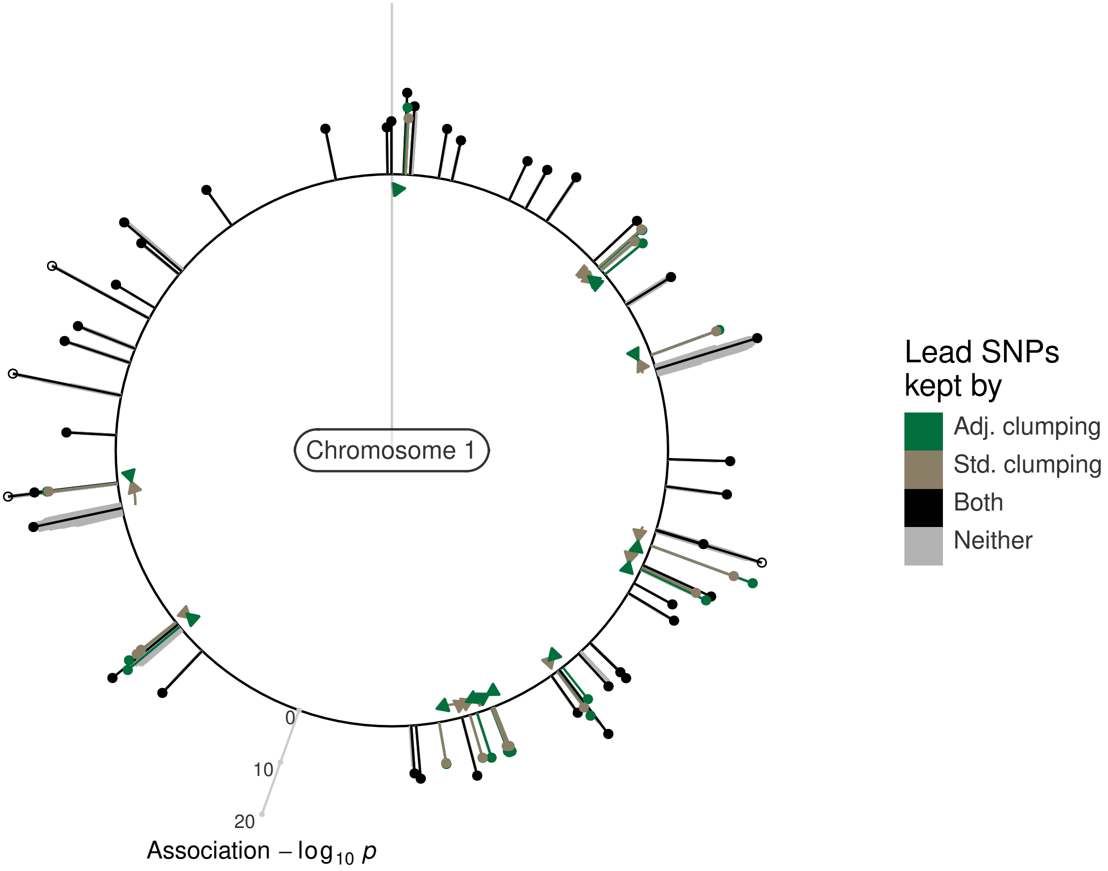


Figure S13: Chromosome 1 association signals for clumping with 5 Mb distances and LD cutoff 0.01. As in the main text, signals are coloured by the LD measure that retains it when clumping. $-\log_{10} p-$values are capped at 20 for open circles. Each SNP kept by exactly one method has an arrow showing the SNP that removed it for the other method. E.g., a green pin shows a SNP kept only with adjusted LD, and has a brown arrow from the SNP that removed it when using standard LD.

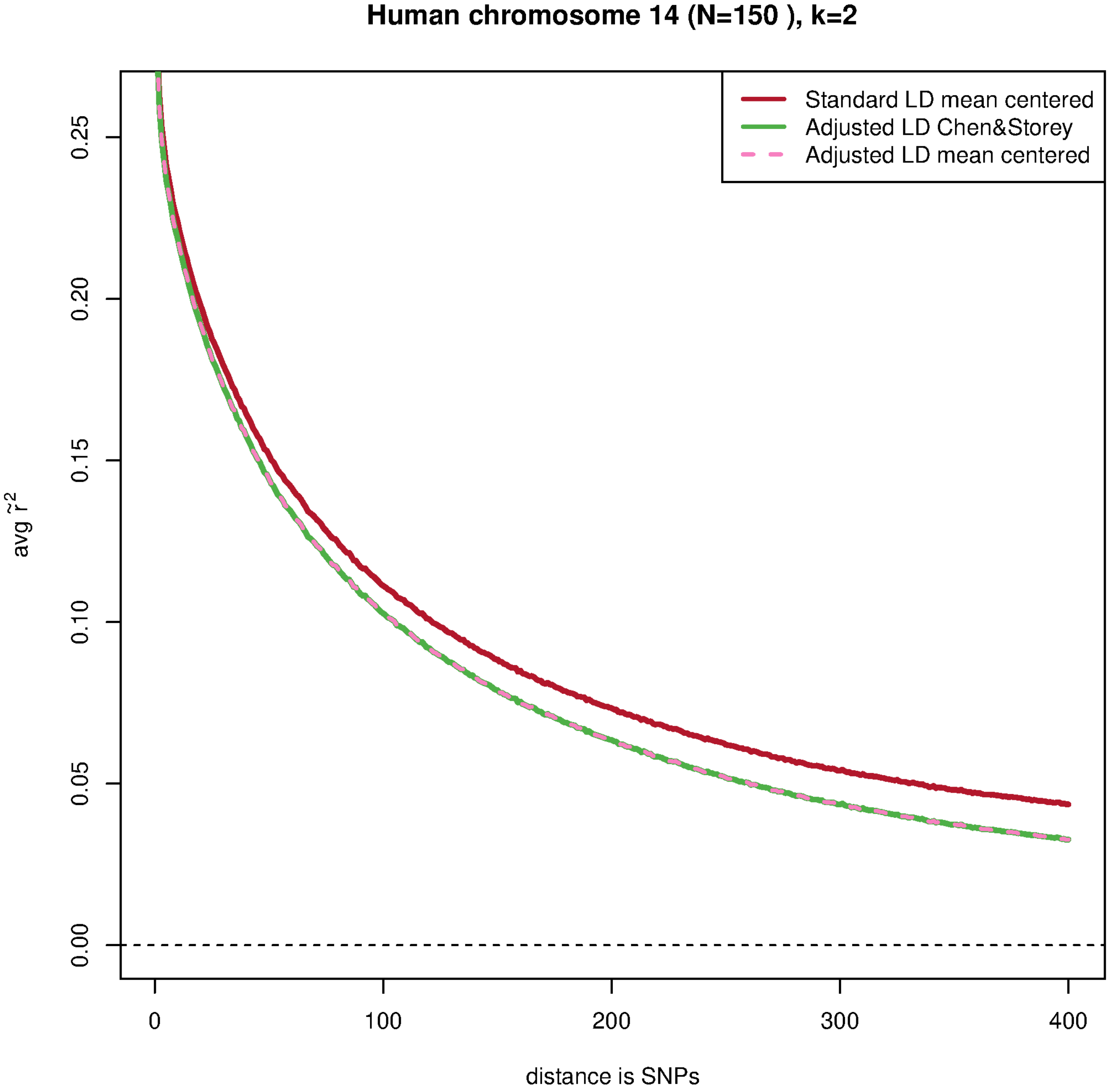


Figure S14: LD decay curves based on the human population (YRI,ASW,CEU) using just the SNPs position rank as distance. We used 3 different approaches for $\tilde{r}^2$. We used both the Chen and Storey [2] and the more standard approach for PCA where the genotypes are just mean centered prior to performing the PCA.

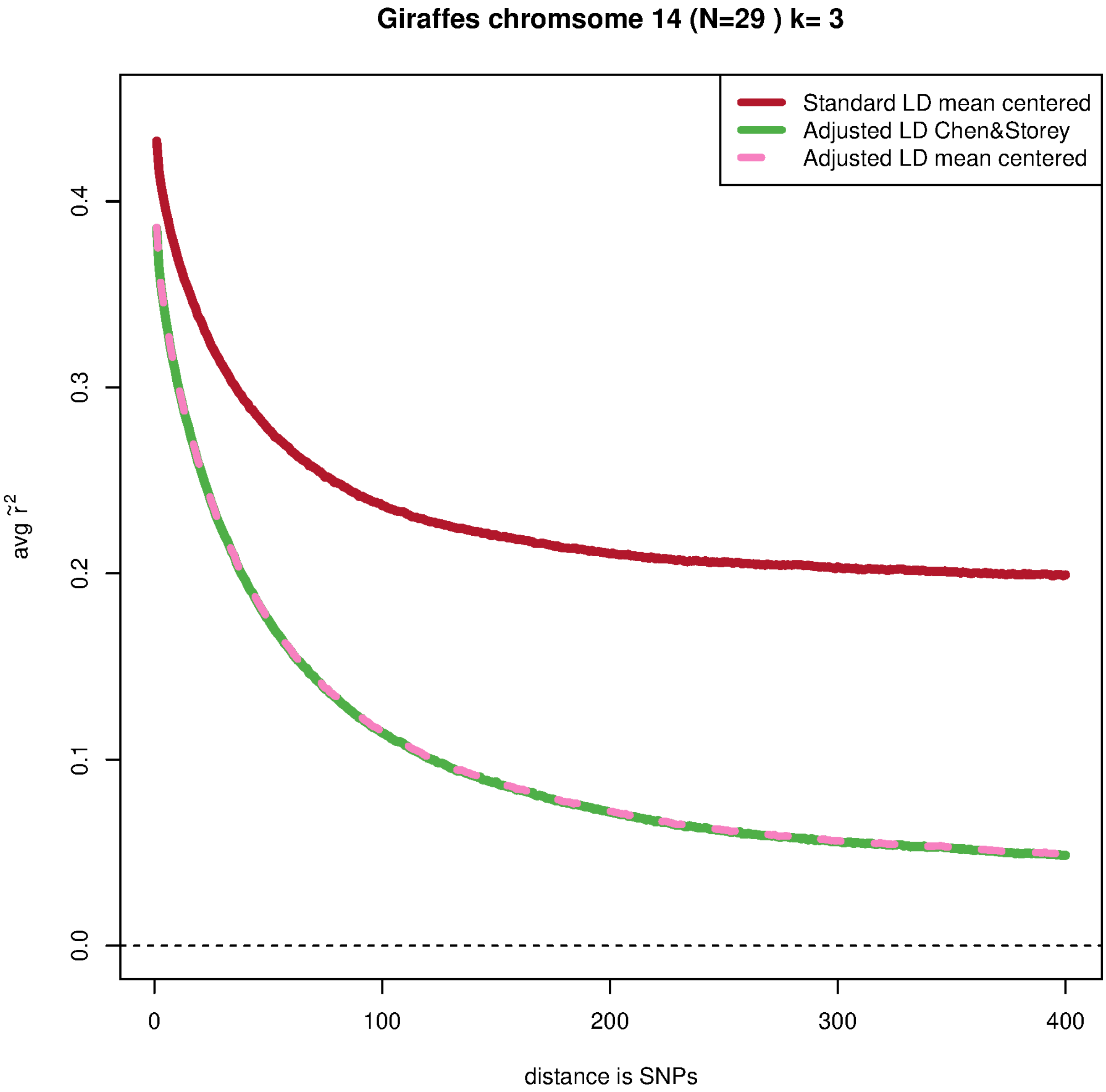


Figure S15: LD decay curves based on the giraffe population (Masai,Reticulated,Southern) using just the SNPs position rank as distance. We used 3 different approaches for $\tilde{r}^2$. We used both the Chen and Storey [2] approach and the more standard approach for PCA where the genotypes are just mean centered prior to performing the PCA.
